# The contribution of an X chromosome QTL to non-Mendelian inheritance and unequal chromosomal segregation in *A. freiburgense*

**DOI:** 10.1101/2023.05.16.540925

**Authors:** Talal Al-Yazeedi, Sally Adams, Sophie Tandonnet, Anisa Turner, Jun Kim, Junho Lee, Andre Pires-daSilva

**Affiliations:** School of Life Sciences, University of Warwick, Coventry, CV4 7AL, UK; Institute of Molecular Biology and Genetics, Seoul National University, Seoul, 08826, South Korea; Liverpool School of Tropical Medicine, Pembroke Place Liverpool, L3 5QA, UK; Department of Convergent Bioscience and Informatics, Chungnam National University, Daejeon, 34134, South Korea

## Abstract

*Auanema freiburgense* is a trioecious nematode with co-existing males, females, and selfing hermaphrodites. Crosses of XO males with XX females result in a low percentage of XO sons due to the elimination of the nullo-X spermatids by the fathers. This process yields most viable sperm containing an X chromosome, leading to a higher transmission probability of the X chromosome compared to expected transmission via random segregation. The mechanism underlying this process involves the asymmetric distribution of essential cellular organelles during sperm formation, which likely depends on the X chromosome. Specifically, sperm components segregate with the X chromosome daughter cell, while other components are discarded in the nullo-X daughter cell. Here we found that intercrossing two strains of *A. freiburgense* results in lines in which males produce viable nullo-X sperm. Thus, crosses of those hybrid males with females result in a high percentage of sons. To uncover the genetic basis of nullo-spermatid elimination and X-chromosome drive, we generated a genome assembly for *A. freiburgense* and genotyped the intercrossed lines. We identified a QTL encompassing several genes on the X chromosome that are associated with its non-Mendelian inheritance observed in *A. freiburgense*. This finding provides valuable clues to the underlying factors involved in asymmetric organelle partitioning during male meiotic division and thus non-Mendelian transmission of the X chromosome and sex ratios.

## Introduction

Animal reproduction involves the production of gametes, with distinct characteristics and patterns of inheritance. Male and female gametes differ not only in size but also in the transmission of cytoplasmic elements. Typically, cytoplasmic components such as mitochondria and bacterial endosymbionts are maternally inherited (but see ^1–4^). In contrast, the nuclear genome is usually inherited equally from both parents, ensuring a balanced contribution of genetic material. However, exceptions to this symmetrical nuclear inheritance exist across various species, ranging from maternal-only to paternal-only inheritance of the nuclear genome (for review, see ^5^). These variations in inheritance patterns for cytoplasmic elements and the nuclear genome highlight the dynamic nature of reproductive strategies in animals. Understanding these exceptions contributes to our broader understanding of reproductive biology and the diverse mechanisms employed by organisms to propagate their genetic material.

One such variation is the asymmetric transmission of sex chromosomes. In XX:XY sex determination systems, for instance, half of the gametes produced by the male have one X chromosome and the other half have one Y chromosome. The transmission is asymmetric because only the male can transmit the Y chromosome to the next generation and females transmit twice the number of X chromosomes. Although the expected XX:XY ratio in the offspring for this type of sex determination is 1:1, selfish genetic elements in one of the sex chromosomes may drive a non-Mendelian transmission, resulting in a bias towards female or male offspring ^6^.

Similarly, XX:XO sex-determining systems may also result in non-Mendelian transmission of the X chromosome, resulting in sex ratio bias after crosses. In this type of sex-determining system, the male is heterogametic and therefore is expected to produce an equal number of X-bearing and nullo-X gametes. In some organisms, however, the nullo-X sperm is not produced by males, resulting in gametes that only carry an X chromosome ^7–9^. Thus, the following generation is biased towards XX animals.

Nematodes are known to have diverse sex determination systems ^10^, which makes them well-suited for studying sex ratios due to their short life cycles, prolific reproduction, and ease of husbandry. In some nematode clades, crosses between XX and XO individuals result in highly biased XX offspring (for review, see ^11^). In only a few cases the mechanisms underlying this sex ratio bias are known. In nematodes that include the genus *Auanema* and *Strongyloides*, for instance, the male-producing nullo-X spermatid is obligatorily eliminated during spermatogenesis ^7, 12–14^.

Nematodes of the genus *Auanema* are trioecious, consisting of XX hermaphrodites, XX females, and XO males. In hermaphrodites and males, an asymmetric division occurs during spermatogenesis, resulting in organelles partitioning to different sides of the dividing cell (Figure 1A) ^12, 13, 15^. During spermatogenesis in *A. freiburgense*, components essential for sperm function, such as mitochondria and organelles containing cytoskeletal proteins necessary for sperm motility, co-segregate with the X chromosome ^16^. Meanwhile, the sister cell lacking an X chromosome and consisting only of autosomes undergoes differentiation into a residual body that is subsequently eliminated together with organelles such as Golgi complex and endoplasmic reticulum ^12, 13, 15^. Consequently, crosses between males and XX females result in the production of mostly XX offspring ^7, 12, 17^.

**Figure 1.**
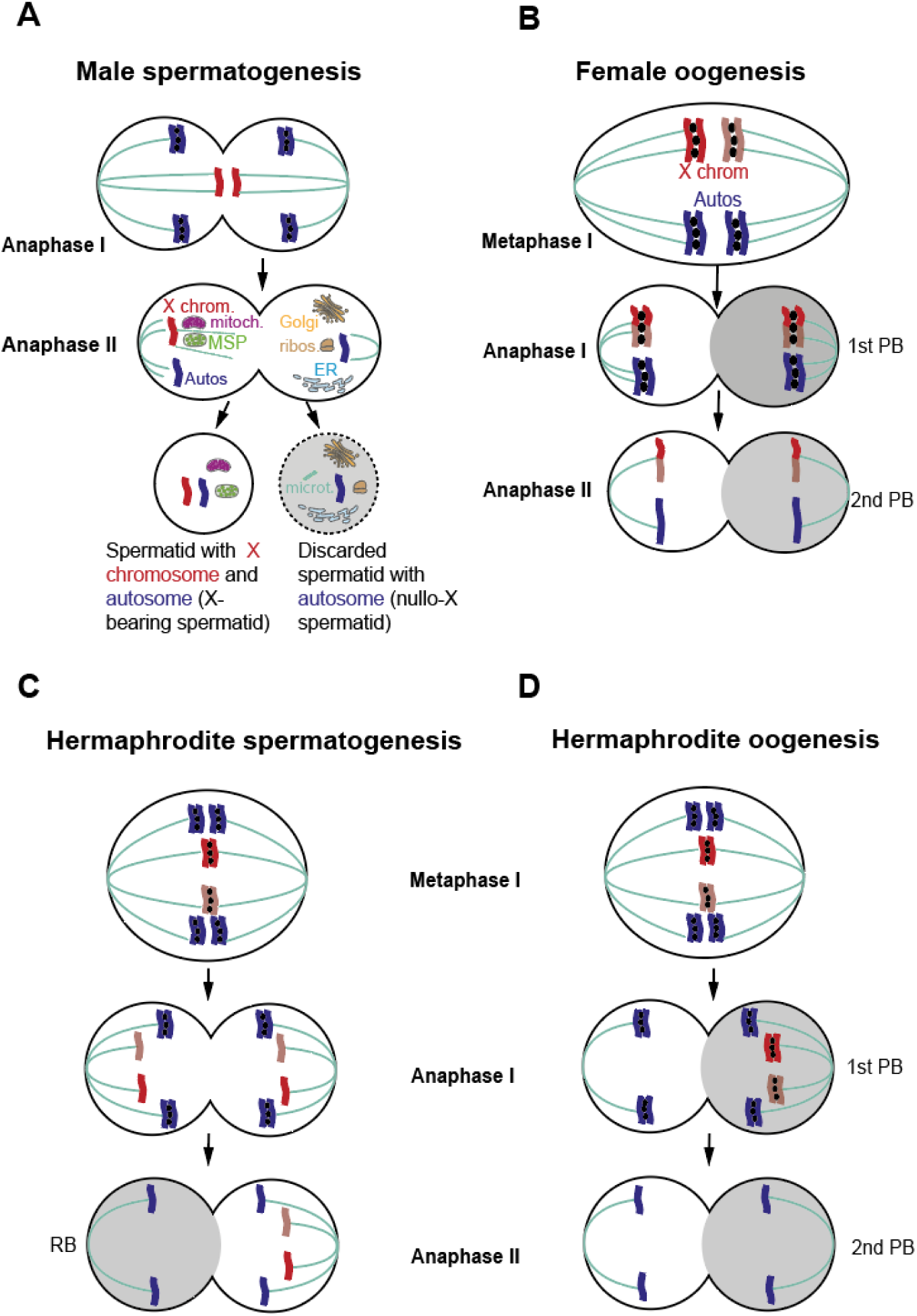
Models of meiosis in *Auanema*. (**A**) In wild-type male spermatogenesis, the chromatids of the X chromosome separate in meiosis I. In anaphase II, the cell that receives the X chromosome also inherits mitochondria and the fibroid bodies containing the Major Sperm Protein (MSP). The cell without the X chromosome (nullo-X cell) inherits non-sperm materials and is discarded as a polar body (not to scale). (**B**) *Auanema* females have canonical meiosis, where the X chromosomes undergo recombination and there is the production of an X-bearing oocyte and three polar bodies. Therefore, most offspring from crosses between females and males are XX. However, if non-disjunction occurs during female oogenesis, it can result in a nullo-X oocyte and male offspring (not pictured in the diagram). (**C**) During hermaphrodite spermatogenesis, diplo-X sperm is produced: the two homologous X chromosomes are segregated into one daughter cell during anaphase II. (**D**) In hermaphrodite oogenesis, the resulting oocyte does not harbor an X chromosome.

Based on the timing of the X chromosome co-segregation relative to the organelles and on the symmetric segregation of organelles in masculinized XX mutants, it has been suggested that the X chromosome may serve as a polarizing signal for this asymmetric cell division ^13, 15^. In this study, we report the use of Recombinant Inbred Advanced Intercross Lines (RIAILs), whole genome sequencing, and Quantitative Trait Locus (QTL) mapping to identify the genetic components controlling asymmetric segregation of organelles in male meiosis in *Auanema freiburgense* ^13, 18^. Some of the RIALs displayed a high rate of male production after outcrossing males. We show that males in these lines produce viable nullo-X sperm, and that this new phenotype maps to a region in the X chromosome. These findings are consistent with the hypothesis that loci in the X chromosome are acting as a polarizing signal during sperm formation, although loci in other chromosomes may also be involved.

## Results

### Auanema freiburgense RIAILs exhibit a transgressive phenotype

*A. freiburgense* is a species with three sexual morphs (XX females, XX hermaphrodites, and XO males) ^18^. Previous research has shown that when an *A. freiburgense* female crosses with a male, the resulting offspring has a skewed sex ratio against males ^13, 18^. An XX sex ratio bias occurs because the X-bearing sperm of the male is viable, whereas the nullo-X spermatid is not ^13^.

Here we tested two *A. freiburgense* inbred strains, APS7 and APS14 ^19^, which produced 18% or fewer sons after outcrossing (Table 1) ^20^.The high ratio of XX offspring confirms previous studies with other strains of the same species ^13, 18^. To investigate sex determination in *A. freiburgense*, we created a genetic linkage map using 100 Recombinant Inbred Advanced Intercross Lines (RIAILs) (Figure 2).

**Figure 2.**
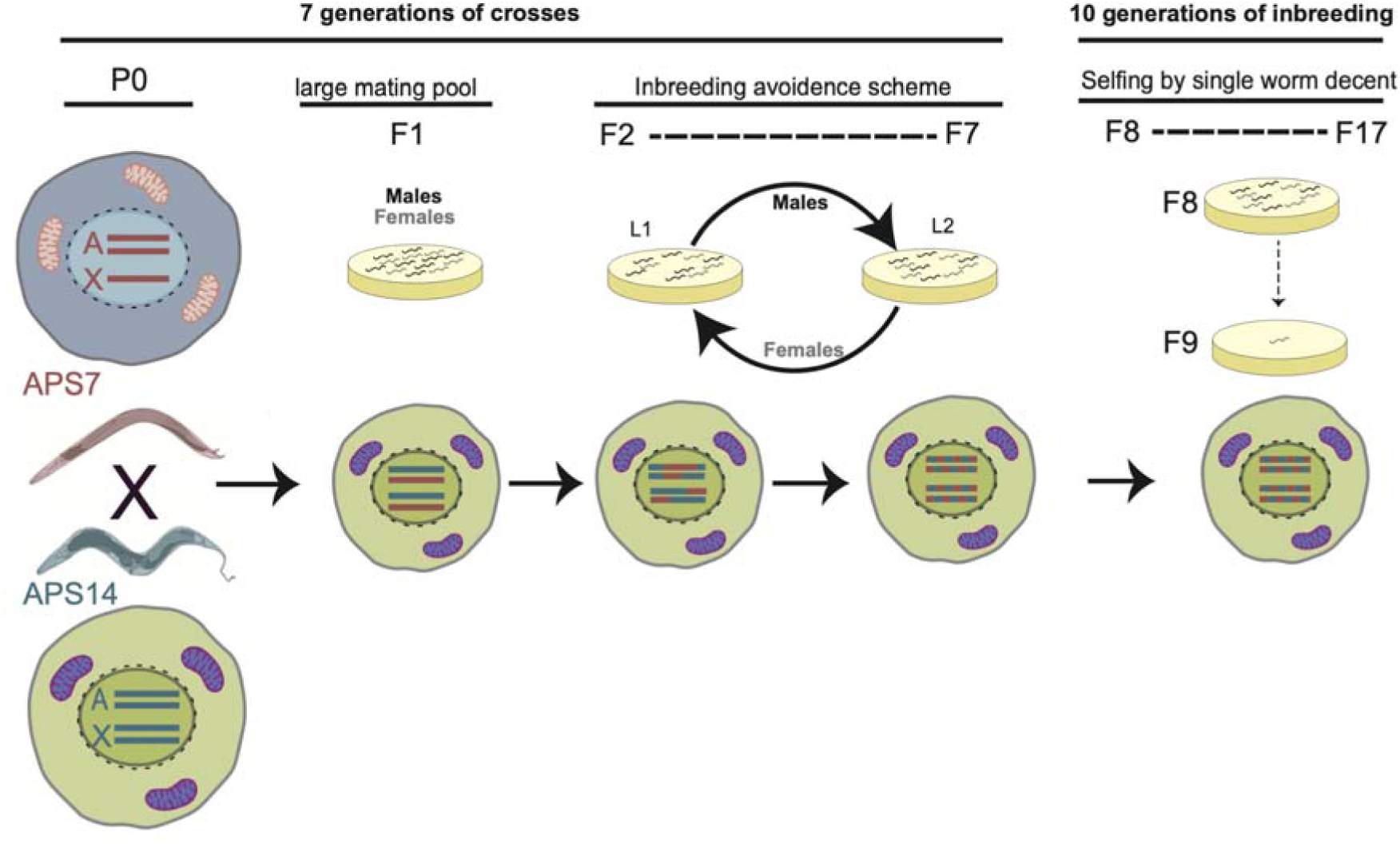
Construction of *A. freiburgense* Recombinant Inbred Advanced Intercross Lines (RIAILs). The breeding scheme involved seven generations of crosses and ten generations of bottlenecking by the propagation of individuals derived from the self-fertilization of a single hermaphrodite. Cells illustrate somatic cells at different stages of RIAILs construction, including mitochondrial and chromosomal background. RIAILs were created by crossing an APS14 female with an APS7 male. F1s resulting from the cross were left to mate in a large mating pool. F1 hermaphrodites and females mated with male siblings were isolated to 100 individual plates to produce F2s, each with unique recombination patterns. From the F2 to the F7 generation, lines were intercrossed in an inbreeding avoidance scheme. In this scheme, each line was crossed with another line only once to maximize haplotype breakpoints. Then, each line was selfed by single worm descent for ten generations to increase genome homozygosity. By the F17 generation, each line had a unique mixed genome from both parental strains and was homozygous at most loci.

**Table 1.**
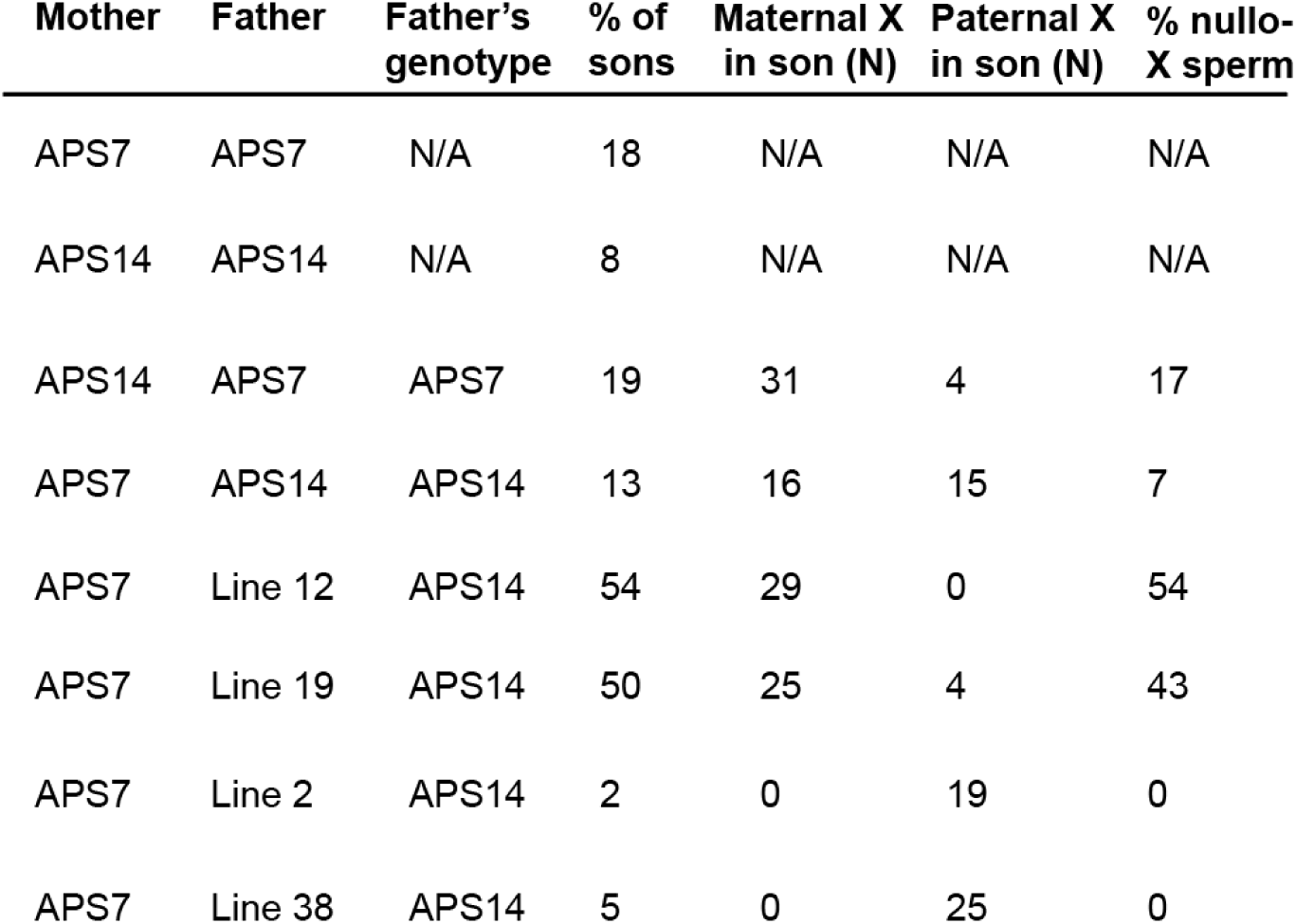
The percentage of male XO progeny and nullo-X sperms from RIAILs, intra- and inter-strain crosses in *A. freiburgense.* The father’s genotype was determined using the X-linked marker X_634_ (see Material and Methods)^20^.

Approximately 60% of these recombinant lines exhibited a transgressive phenotype, indicating the presence of traits not observed in either parental strain. When males from these hybrid lines crossed with APS7 females, they produced a higher percentage of sons (High-male RIAILs, or HM-RIAILs) than the parental strains (Figure 3) ^20^.

**Figure 3.**
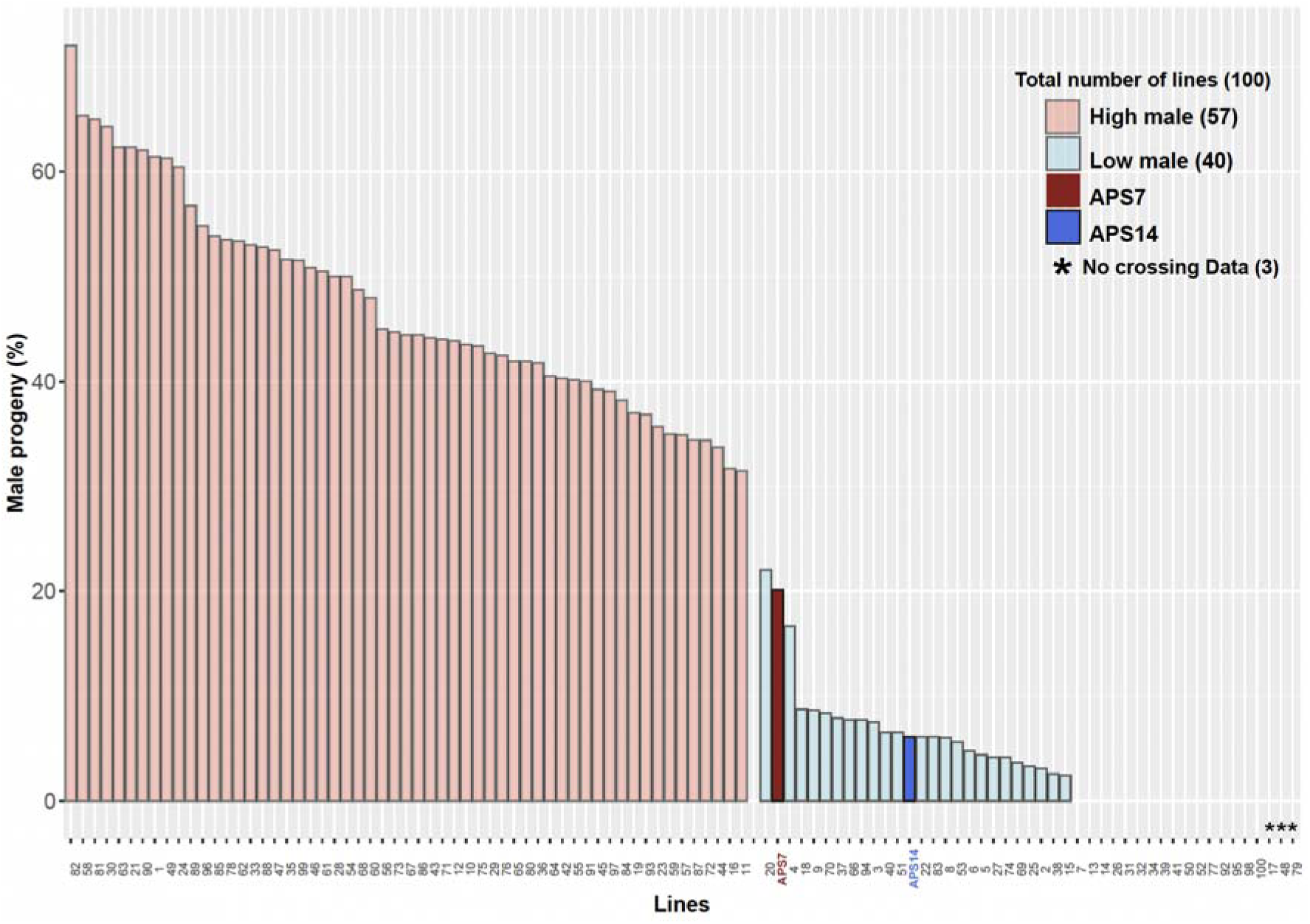
Percentage of male offspring from RIAILs crosses with wildtype APS7 female. In an outcross with a wild-type female, males from most RIAILs produce more sons than the APS7 and APS14 strains. * Indicates that crossing data is not available.

### HM lines produce more viable nullo-X sperm than the parental and LM lines

To monitor the inheritance of the X chromosome after a cross, we used a polymorphic marker on the X chromosome. We found that females and males from APS7 and APS14 produce rare viable nullo-X oocytes and rare viable nullo-X sperm, respectively (Table 1). For instance, in reciprocal crosses between APS7 and APS14, sometimes the sons inherited the X from the mother (indicating fertilization between an X-bearing oocyte and nullo-X sperm), and sometimes from the father (fertilization between a nullo-X oocyte and X-bearing sperm) (Table 1). Overall, the percentage of viable nullo-X sperm produced by APS7 and APS14 males is relatively low (<20%).

Given that the high-male lines described above produce a large percentage of sons, we hypothesized that males in those strains were producing more nullo-X sperm than the parental lines. To explore this further, we focused on two high-male (HM) lines and two low-male (LM) lines. Males from these lines carried the APS14 X genotyping marker and were crossed with an APS7 mother to enable tracking of the X chromosome. Males from the HM lines (lines 12 and 19) produced around 50% of sons, whereas males from the LM lines (lines 2 and 38) generated 5% or less of sons after outcrossing (Table 1). Sons from the LM males inherited the paternal X chromosome in all samples tested, indicating that these lines produce viable X-bearing sperm and few or no nullo-X sperm. In contrast, almost all the sons from the HM males inherited the maternal X chromosome, indicating that ∼50% of the sperm in these lines is composed of viable nullo-X sperm.

### X-chromosome independent high-density genetic linkage map improved X chromosome assembly

To map the locus involved in the production of sons after crossing, the genome of *A. freiburgense* (strain APS7) was sequenced using long reads of Pacific Biosciences (PacBio) and short Illumina pair-end and mate-pair data. The initial genome assembly with this sequencing data resulted in 75 scaffolds (Afr-genome-v1). To improve the genome assembly, the scaffolds were anchored and ordered onto a genetic linkage map. To create this map, we used sequencing data from 100 RIAILs, the parental lines APS7 and APS14, and using Afr-genome-v1 as a reference.

In the related species *A. rhodense,* the X chromosomes do not recombine in hermaphrodites ^7^. Therefore, we expected that the bottlenecking of the RIAILs by selfing single hermaphrodites would still result in an X chromosome containing significant heterozygosity. To accommodate this unique biology, we employed three different approaches to building the genetic linkage map, with parameter settings as follows (see Materials and Methods): (1) using only homozygous markers for all chromosomes (Figure S1A and S2). The resulting map consisted of seven linkage groups, with the X linkage group containing fewer markers than the autosomes (Table S1, Table 2); (2) using both homozygous and heterozygous markers for all chromosomes. However, the markers clustered in several groups (more than seven) (data not shown); (3) a map was constructed independently for the autosomes and another one for the X chromosome (Table 2, Figure S1B and Figure S2). For this latest approach, we used only homozygous markers to group the autosomal markers and both homozygous and heterozygous markers to group the X chromosome markers. Using this approach, the final length of the X chromosome doubled compared to the first approach (Table 2 and Table S1).

**Table 2.** Characteristics of the *A. freiburgense* genome.

| **LG** | **N**<br>**markers** | **Length**<br>**(Mb)** | **N**<br>**genes** | **% repeat**<br>**DNA** |
| --- | --- | --- | --- | --- |
| L.1 | 1079 | 7.2 | 1112 | 19.7 |
| L.2 | 4404 | 13.0 | 1964 | 20.4 |
| L.3 | 3337 | 11.4 | 1833 | 13.6 |
| L.4 | 1638 | 5.3 | 896 | 14.9 |
| L.5 | 2290 | 8.0 | 1442 | 10.9 |
| L.6 | 1412 | 5.0 | 979 | 8.8 |
| L.7 | 2632 | 3.3 | 496 | 35.5 |
| Total | 16792 | 53.5 | 8759 | 16.7 |

When the coding genes from this integrated map-scaffold genome were compared to those of *C. elegans*, a high degree of synteny was observed between L.7 and the *C. elegans* X chromosome (Figure S5). The final genetic linkage map consists of seven linkage groups with 16,792 markers from 29 scaffolds, accounting for 97% of the sequenced genome (Figure 5, Figure S2) ^20^. The overall percentage of missing markers in the GLM is 6% (Figure 5 and Figure S3). Around 3% of sequenced DNA (1,689,119 bp in 47 scaffolds) were left unplaced, as they did not cluster with any linkage group (Table S2). 15 kb of those unplaced sequences correspond to the mitochondrial genome.

**Figure 4.**
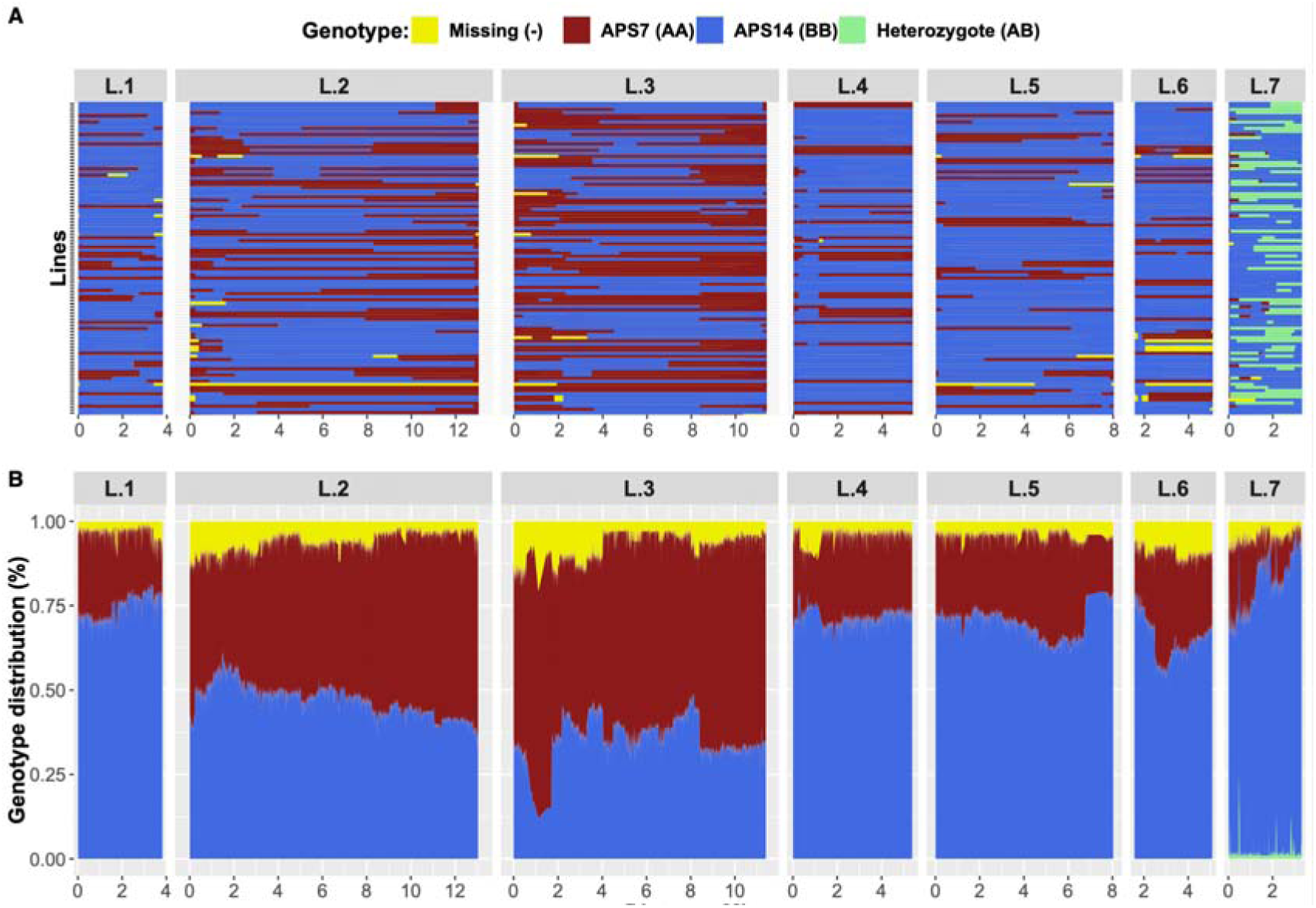
Parental background of the RIAILs, genotype distribution, and recombination across the genetic linkage map. Genotypes clustered into seven linkage groups, with each group appearing in a separate panel on the x-axis. (**A**) Each row in the y-axis represents the genotype of a RIAIL, color-coded according to its genetic background. The x-axis shows the genetic distance in Mb along each linkage group. (**B**) Genome-wide parental genotype distribution, with the color indicating the proportion of parental genotype occurrence (on the y-axis) per genetic distance (on the x-axis). Linkage groups are in separate panels, and the X chromosome (L.7) was plotted separately to show the distribution of heterozygous genotypes compared to homozygous genotypes. Parental genotypes vary consistently per linkage group due to differences in recombination frequencies between linkage groups and within a linkage group. For details about the genotype composition and genotype versus the distance per phenotype of individual RIAILs, see Supplementary Figures S3 and S4.

### Characteristics of the A. freiburgense genome

The resulting chromosomal-scale assembly, which was the result of the integration of the genetic and physical maps, spanned 53.5 Mb (Table 2 and Table S2). The genome contained 89.2% of complete nematode BUSCO orthogroups (Table S3). The annotation of repeat sequences (as described in the Methods section) revealed that 10 Mb (18.2%) of the genome was repetitive (Figure 6 and listed in Table S4). Using a combination of *ab initio* predictors and evidence-based methods (including curated proteins and transcriptomic data), we predicted 8,759 protein-coding genes (Figure 6, Table 2). To assess the completeness and quality of the predicted protein-coding genes, we used BUSCO (nematoda_odb10 database) on the annotated transcript dataset. The annotated transcripts contained 84.8% complete BUSCO orthogroups with 1.5% duplicated BUSCOs. The total percentage of repeat DNA in the table excludes the percentage of repeat DNA in unplaced scaffolds (1.5%).

**Figure 5.**
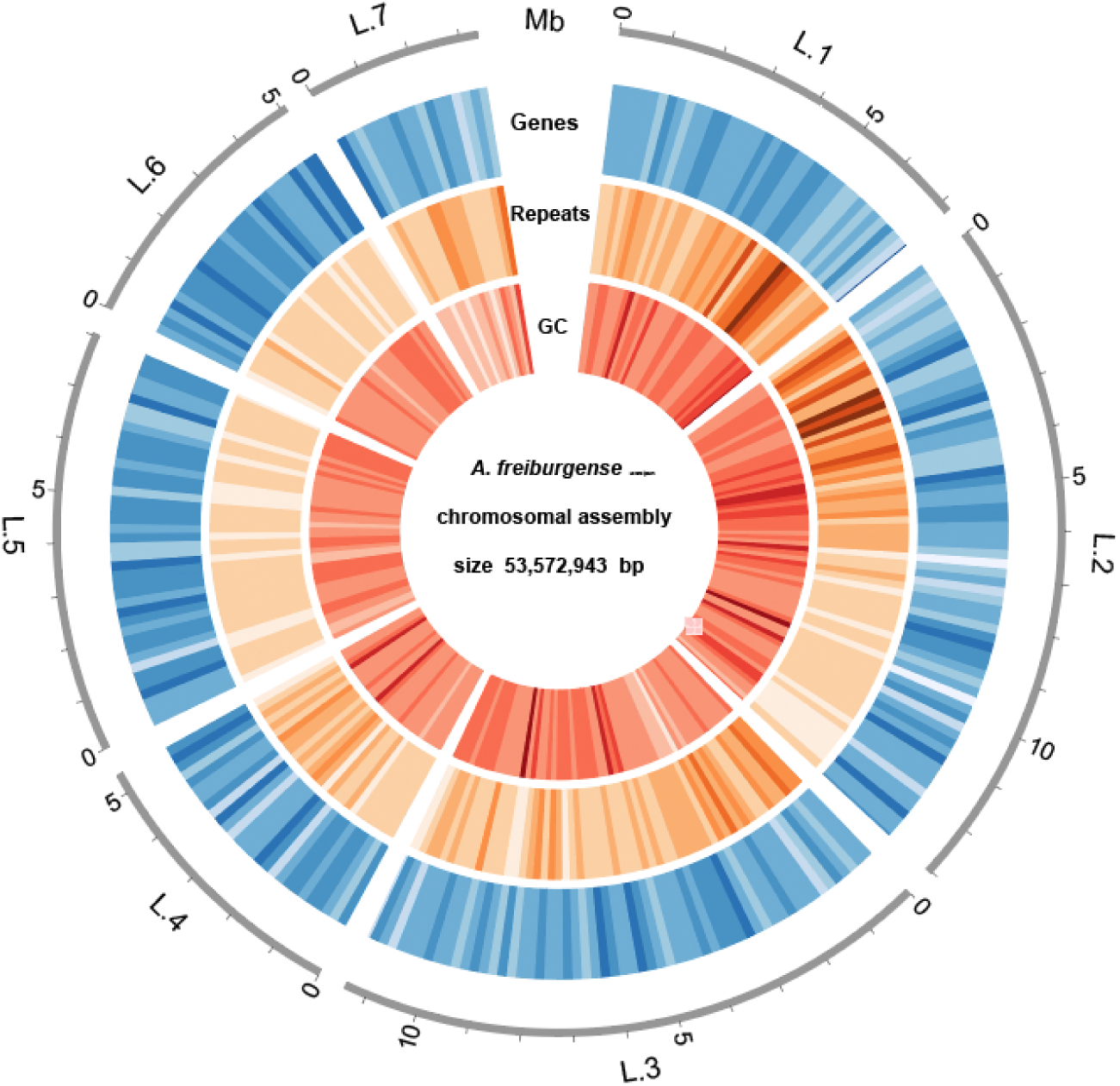
The density of genes, repeats, and GC content across the seven chromosomal-scale scaffolds of *A. freiburgense*. Gene density (blue), repeats (orange) and GC content (red) were plotted along the *A. freiburgense* chromosomal-scale assembly using a window size of 200 kb.

Next, we examined the chromosomal-scale genome of *A. freiburgense* to examine the evolution of chromosomal gene content in nematodes. Previous research has identified seven possible ancestral chromosomal units of Rhabditida nematodes, called Nigon elements ^21, 22^. They represent the reconstructed genic content of the hypothesized seven chromosomes of the ancestor of all Rhabditida nematodes. The Nigon elements in *A. freiburgense* are mostly fragmented into different chromosomes (Figure 7). Notably, Nigon B is fragmented into three different chromosomes in *A. freiburgense* (although it is intact in *A. rhodense*) (Fig 7A). In contrast, Nigon N, which is present in two chromosomes in *A. rhodense*, is mainly in one chromosome in *A. freiburgense* (Figure 7B). When analyzed in context with other nematodes, *A. freiburgense* chromosomes underwent recent fission and fusion events of Nigon elements (Figure 7B, Figure S5, Table S5).

**Figure 6.**
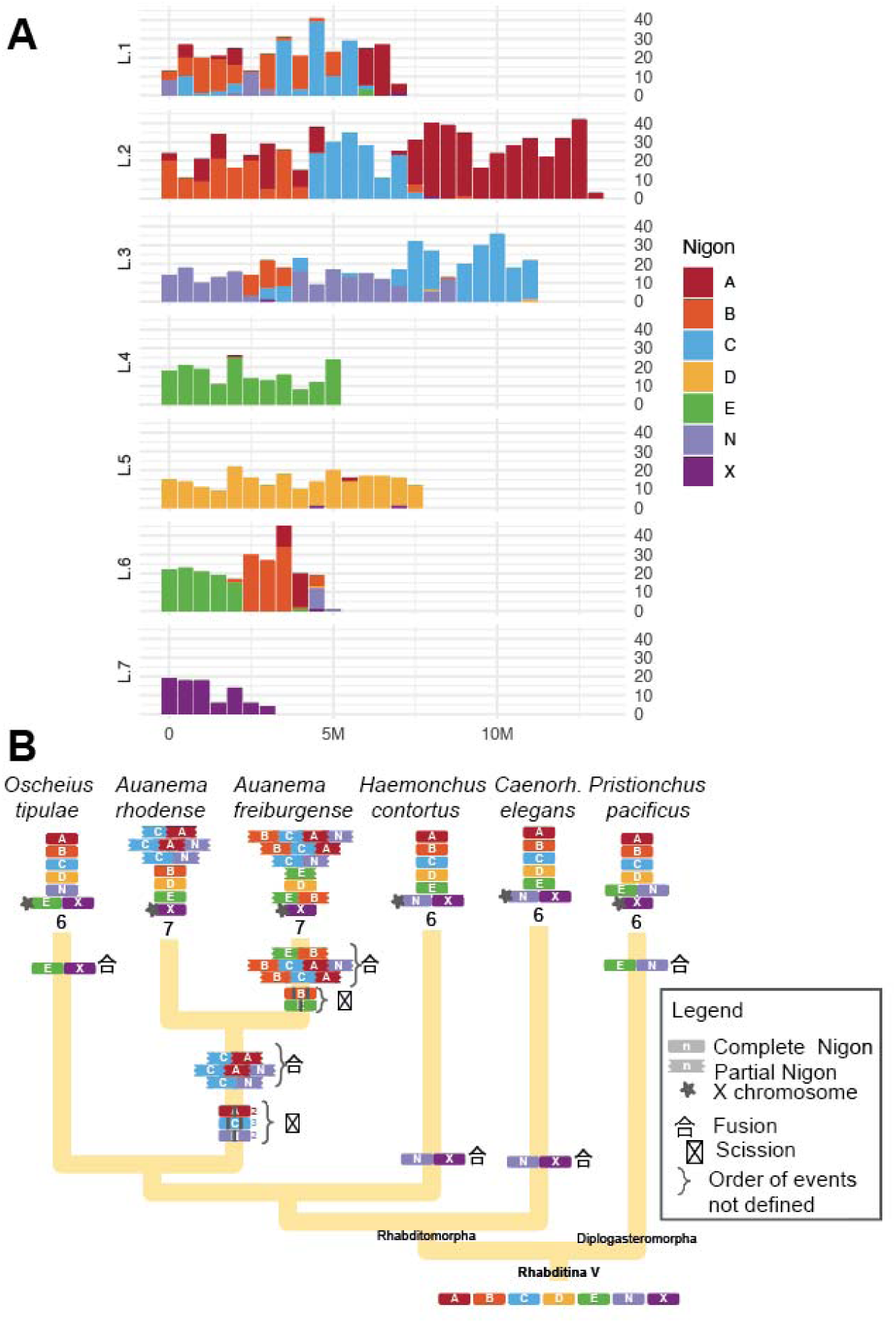
Gene painting according to their Nigon Element association. (**A**) Nigon elements in *A. freiburgense*. **(B)** Model of chromosome evolution in Rhabditida.

### Identifying a candidate region on the X chromosome for the rates of production of males

Next-generation sequencing combined with bulk segregant analysis (NGS-BSA) is a high throughput strategy to identify potential QTL regions associated with traits of interest ^23^. We used NGS-BSA as a first approach, by preparing four pools of DNA, two for HM lines and two for LM lines, each containing DNA from 10 different RIAILs (see Materials and Methods). The delta SNP index refers to the ratio of the frequency of a specific single nucleotide polymorphism (SNP) in a pooled sample of individuals with a particular phenotype (e.g., high-male or low-male) to the frequency of the same SNP in a pooled sample of individuals with the opposite phenotype ^24^. In this case, the SNP index represents the ratio of the frequency of the alternative allele (APS14) to the frequency of the reference allele (APS7). It is used as a measure of the degree of association between a particular SNP and a trait of interest in bulk segregant analysis. After sequencing the four pools, we found two peaks of high delta SNP-index associated with high male rates (Figure 8 and S6). The first region is on L.5, about 6.3 Mb long, and the second is on L.7 (ChrX), approximately 1.0 Mb long (Figure 8, Table S6).

**Figure 7.**
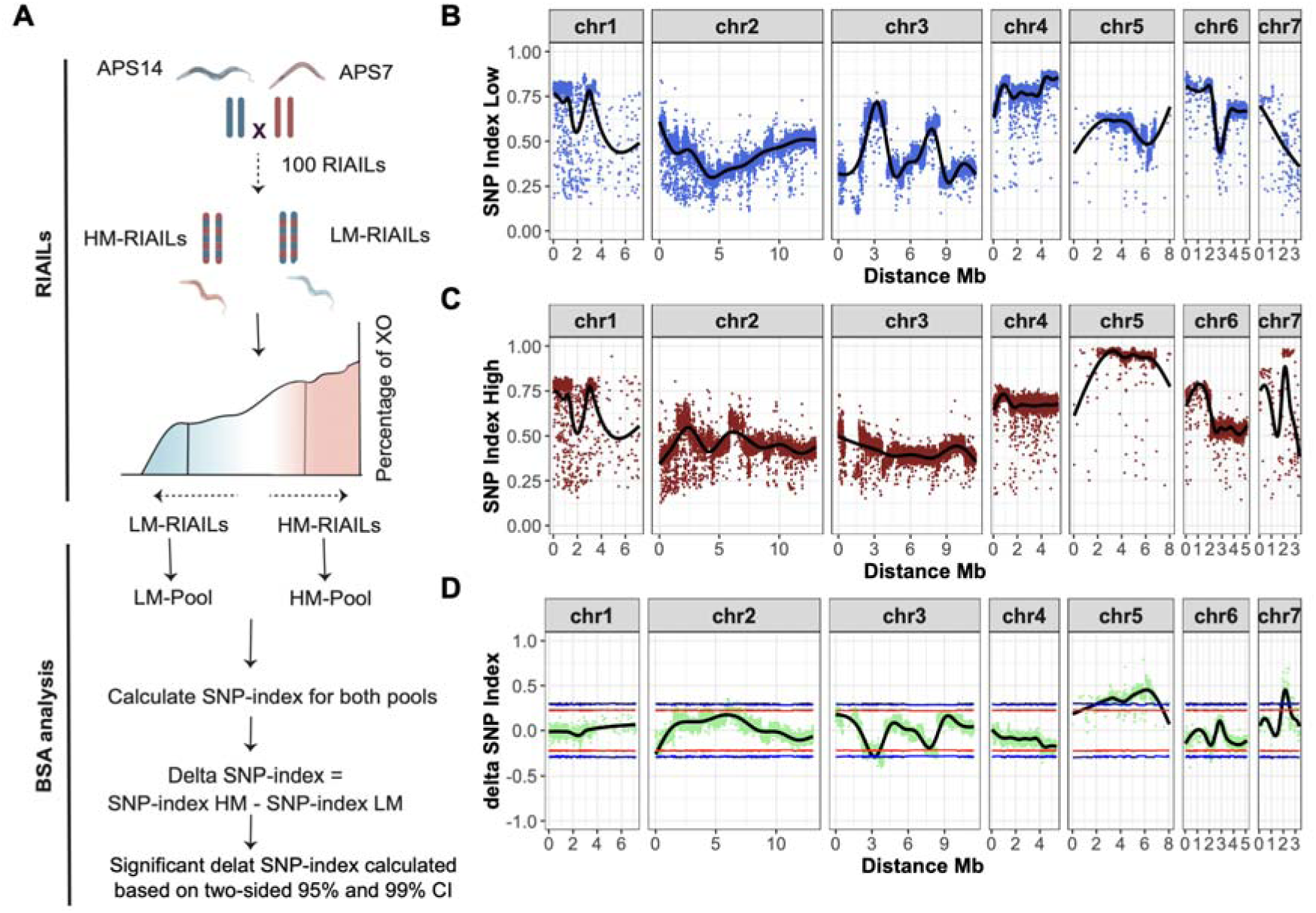
Next-generation sequencing coupled with bulk segregant analysis (NGS-BSA). (**A**) RIAILs were classified as either HM or LM based on the ratio of male progeny resulting from crosses with APS7 females. Variants were identified in two pools of 20 individuals each from each category. The SNP-index was calculated for each SNP from LM-pool (**B**), HM-pool (**C**), and the difference in SNP-index (delta SNP-index) between the two pools (**D**, green dots). The black line represents a tricube-smoothed SNP-index in a sliding window of 1 Mb. The confidence intervals for delta-SNP-index are indicated in red (95%) and blue (99%).

To narrow the candidate locus further, we performed QTL analysis using genome-wide SNPs from individual RIAILs. With this approach, we confirmed the QTL on the X chromosome associated with male rates (Figure 9). The QTL region above the genome-wide threshold at p-value 0.2 spans 0.85 Mb and is located between the positions 1,949,677 bp and 2,806,499 bp on the X chromosome (Figure 10). This region overlaps with the region in the X chromosome identified in the NGS-BSA analysis (Table S6). The peak of the QTL region was determined using the genome-wide threshold at p-value 0.05, which extends from 1,949,688 bp to 2,372,281 bp, with the highest LOD score of 3.37 at position 2,356,542 (Figure 9 ^20^.

**Figure 8.**
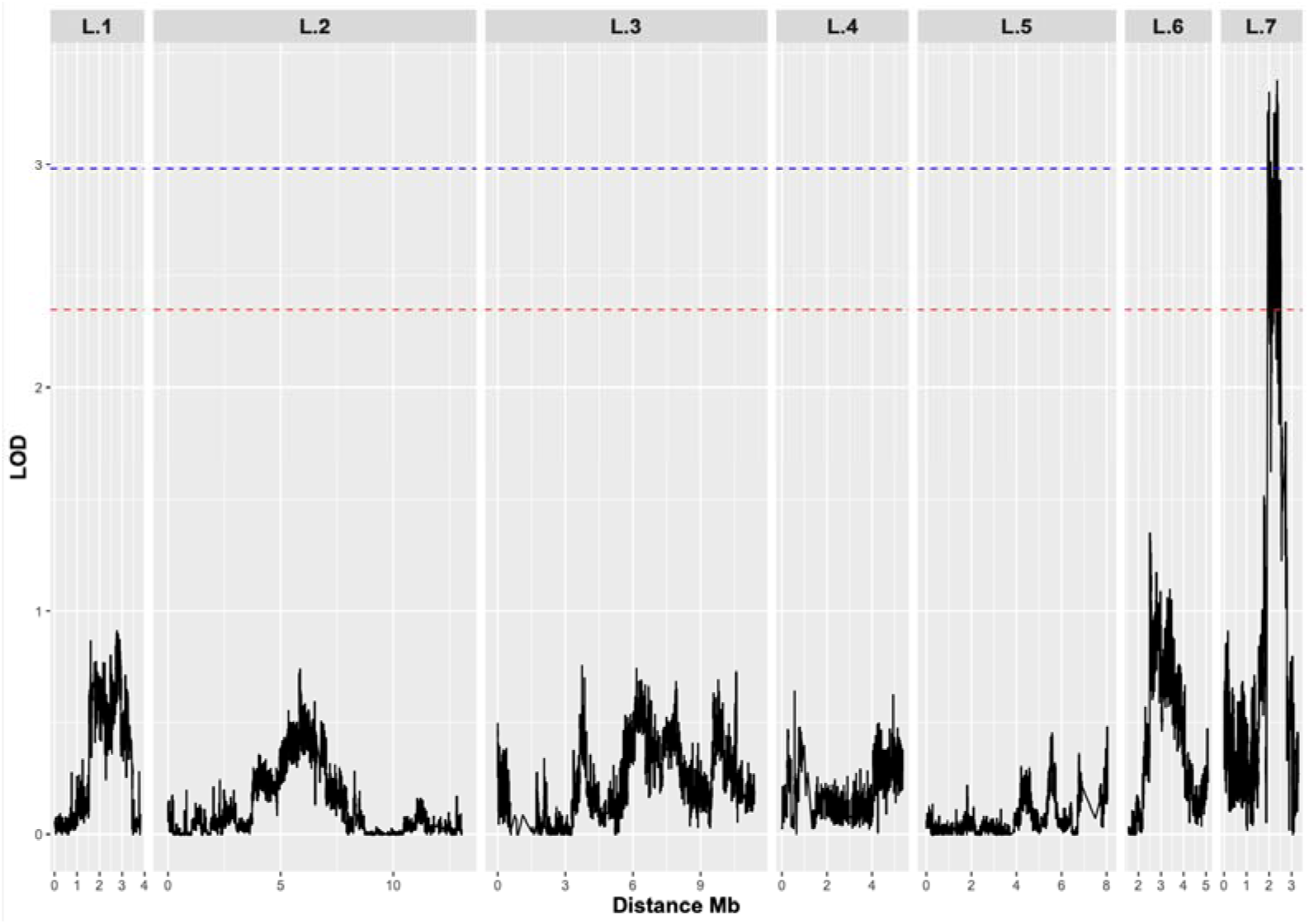
Genome-wide QTL association with the male rate phenotype. The threshold was obtained based on a significance level (p-value) of <0.05 and <0.2, corresponding to LOD (logarithm of odds) scores of 2.98 (blue line) and 2.35 (red line), respectively.

**Figure 9.**
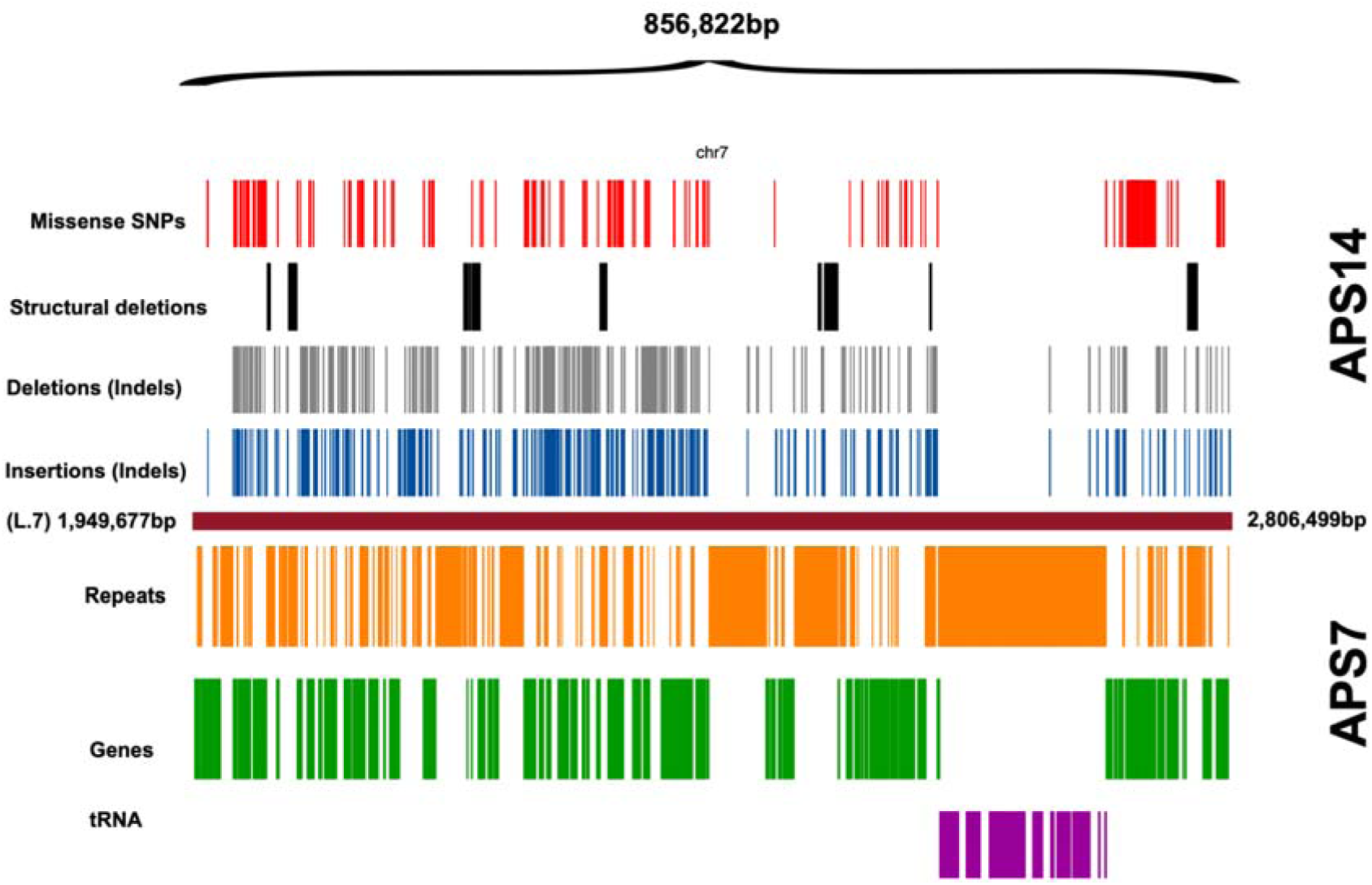
Variants between APS7 and APS14 in the QTL region. The top half of the plot shows the APS14 variants (relative to the reference genome of APS7) with missense variants, large structural variant deletions, and small indels. The bottom half of the plot shows the APS7 annotation of repeats, genes, and tRNAs.

**Figure 10.**
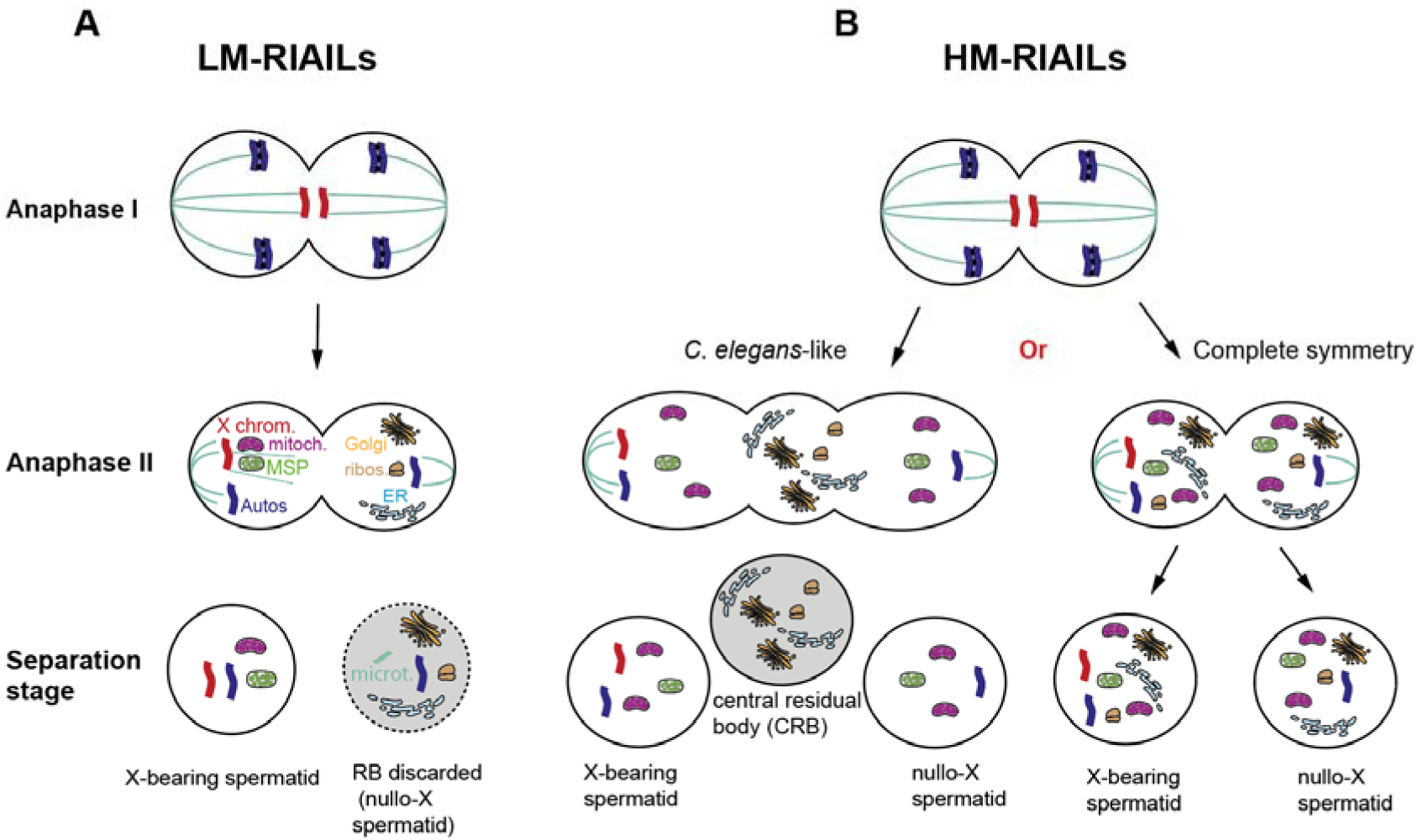
Proposed models of spermatogenesis in HM and LM RIAILs. **(A)** Male cross-progeny resulting from a cross between APS7 females and LM lines inherited the paternal X chromosome, indicating that nullo-X sperm are discarded during the spermatogenesis of males from those lines. **(B)** Nullo-X sperm could be produced by symmetrical distribution of sperm components between X-bearing and nullo-X sperm during anaphase II. The symmetric segregation could be total, where subcellular compartments, including sperm components, segregate equally, or a *C. elegans*-like anaphase II, where a central residual body is formed for discarded materials.

### The QTL region contains significant variation between APS7 and APS14

The variant calling analysis ^25^ detected 354,705 variants between APS7 and APS14, resulting in an average density of one variant per 155 bp. Most of these variants (84.1%) were single nucleotide polymorphisms (SNPs) (Table S7). Of the identified SNPs, 0.1% were nonsense, and 5% were missense SNPs. In addition to SNPs, 8.6% of all variants are insertions, and 7.2% are deletions (Table S7).

The QTL region contains 122 predicted protein-coding genes, 367 repeat motifs, and 307 tRNA sequences ^20^. SnpEff is a variant annotation and effect prediction tool to annotate and predict the effects of genetic variants, such as single nucleotide polymorphisms (SNPs), insertions, deletions, and structural variants, on genes and proteins ^26^. Using this tool, 362 missense variants were predicted within the QTL region on the X chromosome ^20^. 73/122 (59.8%) of the predicted protein-coding genes have at least one missense variation, causing an amino acid change ^20^. In addition, we found 390 insertions, 353 small deletions, and 12 large structural deletions ^20^. The tRNA sequences, specific to APS7, clustered in a region free of protein-coding genes (2,566,140 bp - 2,703,460 bp) (Figure 10) ^20^. Due to the high number of variants within the QTL region, we were unable to pinpoint a particular functional component that regulates organelle distribution during spermatogenesis in *A. freiburgense*. Further testing with functional studies of candidate genes will be required to identify the locus associated with the rates of male production.

## Discussion

*Auanema* populations consist of selfing hermaphrodites, females, and males ^17, 18^. Female versus hermaphrodite development is determined intergenerationally according to maternal age or exposure to social cues ^19, 22, 27–31^, while males are determined chromosomally ^12^.

In *Auanema*, the process of spermatogenesis in XO males and XX hermaphrodites involves the elimination of nullo-X spermatids ^7^. Males produce predominantly X- bearing sperm ^12, 13^, resulting in non-Mendelian transmission of the X chromosome, resulting in mostly XX offspring when mating with females ^17^. When mating with hermaphrodites, males predominantly sire sons ^7^. This occurs due to meiotic non- disjunction of the X chromosome in hermaphrodite oocytes ^7^, which leads to the production of nullo-X oocytes. As a result, the resulting embryos lack an X chromosome, leading to the development of male individuals when fertilized by the X-bearing sperm of the male ^7^.

Unlike in *C. elegans* spermatogenesis, where non-sperm components are deposited into a residual body without DNA ^16^, *Auanema* eliminates those cytoplasmic components together with autosomes. Similar phenomena are observed in other organisms, such as sciarid flies, which undergo asymmetric segregation of chromosomes during meiosis and eliminate an entire set of chromosomes into a residual body (for review, see ^32, 33^). Scale insects also remove sets of heterochromatic chromosomes during spermatogenesis ^34^.

The removal of DNA observed in *Auanema* and other organisms represents a type of Programmed DNA Elimination (PDE), which has independently evolved in different taxa ^35–38^. PDE processes may influence sex ratios in various species. In *Nasonia* wasps, for instance, PDE removes sets of chromosomes to convert female embryos into males ^39, 40^. In the nematode *Strongyloides*, nullo-X sperm, responsible for male development, is eliminated during spermatogenesis ^14^, while in XX embryos, specific portions of the X chromosome are targeted for degradation to generate males ^41^.

Hermaphrodites of the nematode *Rhabdias* also expel one of their X chromosomes into a residual body during spermatogenesis ^42^. The mechanisms of how specific chromosomes are targeted for PDE are still not well understood.

The study of meiotic drive and PDE mechanisms has often been hindered by the lack of genetic tools available for many organisms that exhibit these processes. However, *Auanema* has emerged as a promising model system for investigating these mechanisms due to its advantageous features, including easy husbandry, short life cycle, large brood sizes, and the recent development of genetic tools and resources ^22, 31, 43, 44^.

In this study, we aimed to improve our understanding mechanisms of X-chromosome drive in *A. freiburgense,* generating genetic resources in the form of a chromosomal-scale assembly of its genome. To enhance the contiguity of the assembly, we constructed independent genetic linkage maps for the autosomes and the X chromosome. By accounting for differences in recombination events between these chromosomes during the inbreeding phase of Recombinant Inbred Advanced Intercross Lines (RIAILs), we were able to improve the contiguity of the X chromosome assembly. Our efforts resulted in a 47.3% increase in the size of the X chromosome compared to the initial genetic linkage map (Table S1). This improved assembly will provide valuable insights into the unique biology and mechanisms underlying PDE in *A. freiburgense*.

We serendipitously discovered that by crossing two inbred strains of *A. freiburgense*, males of some hybrid lines can produce more sons than the parental strains. By using genetic markers specific to the X chromosome, we found that males in these transgressive lines retain the nullo-X sperm, which is absent in the parental lines. As a result, when these males mate with females, there is an increased proportion of sons. This ‘new phenotype’ is typical of transgression, which is defined as the appearance of a phenotype in F2s that is more extreme than the parental phenotypes ^45–48^.

Novel allele combinations can result in phenotypes that exceed the range observed in either parent, often with extreme or unexpected traits ^47^. However, this mechanism is unlikely to be the case for high male production in *A. freiburgense*, given that all lines with transgressive phenotypes are inbred and therefore likely to be homozygous in most loci. Furthermore, the parental inbred strains APS7 and APS14, which were originally derived from the outbred strains SB372 ^18, 49^ and JU1782 ^29, 50^ respectively, did not show transgressive phenotypes during the process of inbreeding.

Lines with high-frequency males had an overrepresentation of homozygous APS14 alleles in a region of the X chromosome, indicating that this region is associated with the phenotype. Since the same allele combination was present in the parental line APS14, we can discard a model based on dominance or exposure of recessive alleles ^47^. Instead, the ‘high male’ phenotype is likely to be the result of epistatic interactions between a combination of homozygous alleles of more than one locus. Although the BSA indicated that a second locus associated with high production of males is located in chromosome 5, we could not confirm this by QTL mapping using whole genome sequencing of all RIAILs. The success and precision of NGS-BSA rely on the high number of individuals used per bulk and the coverage of sequencing. Even though we sequenced bulks at high coverage, there were only 10 lines per bulk. We reason that the genetic variation in the small number of lines selected per bulk did not represent the genetic variation in all of the RIAILs sharing a similar phenotype. As a result, another candidate region that is probably not associated with the phenotype was detected. To improve confidence in NGS-BSA, future experiments should increase the number of lines per bulk by pooling all the HM-RIAILs in one bulk and the LM-RIAILs in another bulk to capture all the genetic variations within the RIAILs.

The nature of nullo-X elimination by a polarizing cue from the X chromosome is still unclear. In the ∼800 kb mapped QTL region there are 123 predicted protein-coding genes, and 75 of the candidate protein-coding genes have at least one missense mutation. In addition, there are structural variants in non-coding regions that may mediate the polarization of cytoplasmic components during meiosis. Further investigation of those variants could shed more light on the biological mechanism underlying DNA elimination during sex determination by polarizing the spermatocyte cytoplasm to generate viable sperm and non-viable spermatids.

## Author contributions

T.A. and A.P.S. designed the study. T.A., S.A., A.T., and J.L. conducted the experimental work. Genome assembly, annotation, and analysis were conducted by T.A., S.T., S.A., and J.K. The article was written by T.A. and A.P.S. with input from all other authors.

## Supporting information

Peer review report

## Acknowledgments

A.P.-d.S. and S.A. were supported by grants from BBSRC (BB/L019884/1) and Leverhulme Trust (RPG-2019-329). S.T. was funded by a full Ph.D. scholarship from the program Ciência sem Fronteiras (CNPq agency, process number 201116/2014- 6). A.T. was funded by the Doctoral Training Program from Natural Environment Research Council (NERC CENTA). J. L. was supported by Samsung Science and Technology Foundation SSTF-BA1501-52.

## Materials and methods

### Strains and their maintenance

*A. freiburgense* (previously known as *Rhabditis* sp. SB372, or *A. freiburgensis*) was first isolated in Freiburg, Germany, by Prof. Walter Sudhaus from a horse dung pile ^18, 49^. The *A. freiburgense* SB372 strain underwent eleven generations of bottlenecking (expansion from a single-selfing hermaphrodite) to produce the inbred strain APS7 ^19^. *A. freiburgense* JU1782 strain was isolated from a rotting *Petasites* stem sampled in Ivry (Val-de-Marne, France) ^30^ by Marie-Anne Felix. JU1782 underwent bottlenecking for ten generations and was renamed inbred strain APS14. Nematodes were maintained on NGM plates seeded with *Escherichia coli* OP50-1 at 20 °C.

### Sexual morph identification and female-male cross

In uncrowded conditions, *A. freiburgense* hermaphrodites produce mostly female and male progeny ^29, 50^. *A. freiburgense* dauer larvae invariantly develop into self-fertilizing hermaphrodite adults ^18, 29^. Therefore, to isolate female and male sexual morphs for crosses, dauer larvae were incubated on NGM OP50-1 plates at a low density (3 dauers per 2 cm diameter OP50-1 bacterial lawn) until they reached adulthood and began laying eggs (approximately 48 h after collection) ^19^. Approximately 36 h after the start of egg-laying, L2 stage female larvae were differentiated from males by tail morphology and moved to fresh female-only plates, to prevent fertilization. After 24 h, virgin females reached adulthood and were used in crosses. Young adult males were isolated from the original low-density plates.

The crosses were conducted with a ratio of one female to one male. The male was subsequently removed after 24 hours, and the mother was transferred to a new plate. The male and non-male progeny were counted 48 to 72 h after egg laying and the counts were combined for all 3 plates for each cross.

### Generation of A. freiburgense Recombinant Inbred Advanced Intercross Lines (RIAILs)

One hundred *A. freiburgense* recombinant inbred advanced intercross lines (RIAILs) were generated by crossing APS7 males with APS14 females. Hybrid F1 progeny resulting from the cross was left to mate (or self-reproduce) in a large mating pool. F1 self-reproducing hermaphrodites and females, each mated with one or more males, were isolated to establish the lines. Three to five F2 females were picked from each line and crossed with two males from a different line, in an inbreeding avoidance scheme. Inter-crosses between lines from F2 to F7 were established in a way that every two lines were only crossed once to maximize haplotype breakpoints. After seven generations of inter-line crosses, lines were inbred by single worm descent for ten generations to bring alleles into homozygosity at most loci. Once the 17th generation was reached, the 100 lines were maintained by sampling animals from a crowded plate to a fresh new plate once a month.

### Crossing males from A. freiburgense Recombinant Inbred Advanced Intercross Lines (RIAILs) with wild-type APS7 females

In each cross, a male from each RIAIL was crossed with a wild-type APS7 female for ∼24h. After the cross, males were removed. The fertilized female was moved to a new plate every day until it stopped laying eggs, to synchronize the growth of the F1 progeny. Ratios of male-to-female progeny were scored for each cross. Males from all lines were crossed except lines number 17, 48, 79, and 107.

### DNA Extraction from A. freiburgense Recombinant Inbred Advanced Inter-crosses Lines (RIAILs) and sequencing

For each line, the nematodes were harvested with water from five NGM plates (10 cm in diameter). They were collected in a conical tube and washed 2-3 times with water. After each wash, nematodes were let to settle naturally to the bottom of the tube rather than by centrifugation. 500 μl of lysis buffer (100 mM Tris (pH 8.5), 100 mM NaCl, 50 mM EDTA, 1% SDS, and 1% beta-Mercaptoethanol) was added and tubes were frozen at -80 °C overnight.

Three cycles of thawing and freezing were performed before adding 2.5 μl of proteinase K (20 mg/ml) to each tube, followed by incubation at 65 °C for 3-4 hours. DNA extraction was performed with the Gentra Puregen Core kit (Qiagen) following the manufacturer’s instructions.

Sequencing libraries were generated using TruSeq DNA nano gel free at the GenePool facility at the University of Edinburgh. Sequencing libraries were sequenced using the Illumina HiSeq platform to generate 150-bp paired-end reads, with an insert size estimation of 350 bp.

### DNA extraction from A. freiburgense for PacBio, Illumina mate-pair, and pair-end sequencing

For PacBio long-read sequencing, DNA was extracted from plates containing *A. freiburgense* APS7 of various stages. To lyse worms, we used 6 mL of 55 °C Cell Lysis Solution from The Gentra Puregene® Cell and Tissue Kit (Qiagen) with 0.1 mg/mL proteinase K and 1% β-mercaptoethanol. The solution was directly poured into a 200-µL worm pellet, and worms were lysed in the solution at 55 °C for 8 h with occasional inverting. Polynucleotides were purified from the mixture by using phenol-chloroform-isoamyl alcohol (25:24:1 v/v) DNA extraction and ethanol precipitation methods coupled with phase-lock gel to minimize pipetting and DNA shearing.

Purified polynucleotides were re-dissolved in the TE buffer and treated with 10 μg/mL RNase for 2 h. DNA was extracted by the same DNA purification procedure, and dissolved in 10 mM Tris-HCl (pH 8.0). Macrogen (South Korea, https://www.macrogen.com/en/main) performed PacBio library preparation and sequencing in the Sequel platform with the continuous long-read sequencing mode.

DNA extraction for Illumina mate pair and pair-end sequencing of APS7 was performed as described for the RIAIL sequencing, using DNA from dauer larvae collected from 200 plates, as previously described ^51^.

### Genome assembly

The assembly of the PacBio data was performed with the Canu software (version 1.6) ^52^, using a minimum read length (minReadLength) of 3 kb and setting the corrected error rate (correctedErrorRate) to 0.030, as this resulted in the most contiguous assembly. The estimated genome size (genomeSize) was set to 55 Mb.

Bacterial contamination was identified and removed through BLASTn (version 2.7.1) ^53^ alignments to a contaminant database that included 3,000 bacterial genomes downloaded from the European Nucleotide Archive (ENA) on March 30, 2018. The following BLAST parameters were used "-task megablast -evalue 1e-06 -outfmt 6 - perc_identity 50".

We first polished our preliminary draft genome using Quiver with PacBio raw reads aligned to the assembly using pbalign (version 0.3.1). We then further polished the assembly using Pilon with all available Illumina short reads. Illumina short reads were first preprocessed using skewer (parameters " -n -Q 20 -l 51"). Trimmed reads were then aligned against the assembly using BWA. These alignments were then used to fix base-level inconsistencies between the Illumina read and the genome assembly using Pilon (“--changes --fix bases --chunksize 8000000 --diploid ^54^.

A list of all the programs used in all bioinformatics analyses, their versions, and parameters are included in Table S10.

### RIAILs genotyping

The assembled genome polished with Illumina reads was used as a reference genome to genotype the 100 RIAILs and the APS14 parent samples. The quality of individual samples’ raw reads was checked using FastQC to ensure sequencing adaptors were removed. Each line/strain paired-end DNA sample was aligned against the long-read genome assembly using BWA aligner ^55^. Sequence Alignment Map (SAM) files were converted into Binary Alignment Map (BAM) files, then sorted according to the position in the reference genome ^56^. Aligned reads in every BAM file were assigned a new read-group tag using the Picard AddOrReplaceReadGroups tool to make it compatible with the genome analysis toolkit GATK pipeline ^57^.

Variants from every alignment bam file with a new read-group tag were identified against the APS7 reference genome using genome analysis toolkit GATK tools, producing a single variant calling file containing the variants of each sample against the reference APS7 genome. Low-quality variants were filtered with VariantFiltration provided by GATK tools using default parameters ^25, 57–59^. Then variants were further filtered using vcftools keeping only variants with a minimum depth of 10, and a minimum genotype quality of 30, removing variants that are missing in more than 25% of RIAILs, removing all the indels, and only keeping bi-allelic SNP variants ^60^. The final 274,394 SNPs were used as markers to construct a genetic linkage map.

### Construction of a high-density genetic linkage map

The genetic linkage map was constructed using R/qtl and ASMap R packages ^61, 62^. R/qtl was used to process markers pre-construction of the genetic linkage map and ASMap for the genetic linkage map. Markers were imported into R using rqtl as RILs data, crosstype = “riself”, expecting no heterozygous markers in the dataset, heterozygous markers were assigned a missing value. 10,326 markers either missing the genotype of one of the parents or found heterozygous were omitted from the data set, leaving 264,068 markers from 100 samples on 40 scaffolds. Using the rqtl function ‘drop.markers’ the marker set was further refined by (i) removing markers that were absent in 90% of the data allowing only 10% of missing values, (ii) removing markers that shared similar genotypes to reduce redundancy in the data set and (iii) removing markers with abnormal segregation distortion (P. value < 0.05). After that, ASMap R package was used to construct a genetic linkage map with the remaining 14,955 markers from 100 samples on 25 scaffolds. The genetic map was constructed using the ‘mstmap.cross’ function provided by the ASMap package using a p value of 1e-11. Genetic distance between markers was computed and markers were linked and organized into 7 main linkage groups. An initial genetic linkage map of seven linkage groups was constructed using 14,884 markers from 100 samples on 22 scaffolds representing 93.7% of the genome.

To improve X chromosome assembly, another genetic linkage map was constructed specifically for the X chromosome by including all the heterozygous and homozygous markers and removing all autosomal scaffolds. Linkage group 7 from the initial genetic linkage map was identified using synteny mapping as the X chromosome in *A. freiburgense*. To construct an independent genetic linkage map for the X chromosome, markers were imported into rqtl as “f2 intercross” to include all the heterozygous and homozygous markers. In total, 264,068 markers from 40 scaffolds were imported. Using the rqtl function ‘drop.markers’, we removed markers that were not present in 75% of the data allowing only 25% of missing values, and redundancy in the data set was reduced by removing markers that share similar genotypes. Then, markers belonging to autosomal linkage groups (L.1 to L.6) were removed leaving only 21 scaffolds with 3,128 markers from X chromosome scaffolds and unplaced scaffolds. Before linkage map construction, using ASMap “pullCross” function, markers with more than 10% missing values and those with a high segregation distortion (segregation ratio less than 1:98:1) were removed to be pushed back into the map after construction. The X chromosome genetic map was constructed using the ‘mstmap.cross’ function provided by the ASMap package using a p value of 1e-12. Using the “pushCross” function most markers with high segregation distortion were pushed back to the constructed map using a segregation ratio threshold of (0.5:99:0.5). Small linkage groups were subsetted leaving only the main linkage group representing the X chromosome with 2,632 markers. The new X chromosome genetic linkage map was added to the previously constructed genetic linkage map replacing Linkage Group 7 (L.7). The final genetic linkage map of 7 linkage groups contained 16,792 markers from 29 scaffolds and represents 97% of the genome ^20^.

### Synteny analysis and identification of ancestral chromosomal elements (Nigon Elements)

Scaffolds were anchored to the genetic map using ALLMAPS software, producing a chromosomal-scale assembly, where scaffolds were ordered and oriented into their respective position within each linkage group ^63^. The chromosomal assembly was aligned to the *C. elegans* genome using MUMmer with default parameters and macro-synteny patterns were visualized using Circos ^64, 65^.

To assess the completeness of the genome assembly, a BUSCO analysis was conducted in the genome mode using Augustus gene discovery against the latest nematode database (nematoda_odb10) (Table S3 and S8). To examine the patterns of Nigon Elements in each chromosomal-scale scaffold, we used the program vis- ALG (https://github.com/pgonzale60/vis_ALG) ^21^. Briefly, we used the location of the BUSCO genes identified in *A. freiburgense* and the association between BUSCO genes and Nigon Elements previously determined to paint the chromosomal-scale scaffolds according to the Nigon Elements.

### X-chromosome genotyping and calculation of nullo-X ratio

To follow the X chromosome inheritance, we used an X-linked polymorphic marker (X_634_) where a HindIII restriction site (AAGCTT) is present in APS7 but not in the APS14 strain (AAACTT). X chromosome genotyping was conducted by PCR amplification of a region spanning the SNP followed by digestion of the product using HindIII. Genomic DNA was extracted from individual males using a modified single-worm PCR method. A single male was frozen in 20 µl of 1x PCR buffer at -80 °C for a minimum of 24 h. After thawing, the tissue was lysed and genomic DNA was released by the addition of 0.5 µl of proteinase K (20 mg/ml) and incubation at 65 °C for 60 mins, followed by 95 °C for 15 mins, to inactivate the enzyme. Samples were frozen at -20 °C for at least 24 h before use in PCR. Each PCR reaction was conducted with 2 µl of DNA, 10 µl of GoTaq Green Master Mix (Promega), 10 µM forward primer (UW634_F 5’-AGGGACACGATTGCCTTCTG-3’), and 10 µM of the reverse primer (UW635_R 5’-AATGCCGCGGAGGTCTTTAA-3’) in a final volume of 20 µl. The following cycling conditions were applied: 94 °C for 5 min, followed by 30 cycles of 94 °C for 15 sec, 55 °C for 30 sec, and 72 °C for 1 min. The PCR products were digested by direct addition of 0.5 µl of HindIII (Promega) and incubation for 1 h at 37 °C. The genotype of each sample was determined by agarose gel electrophoresis of the digested sample. The APS7 X allele gave 2 fragments (329 bp and 234 bp) and the APS14 allele remained undigested (563 bp). Note the X_634_ marker region is outside the QTL region (coordinates 438,785 to 439,347).

To determine the percentage of nullo-X sperm that contributed to the generation of sons, the paternal X chromosome was genotyped using the X-linked marker X_634_ (Table 1). To determine the percentage of viable nullo-X sperm for each strain, we multiplied the percentage of sons resulting from crosses by the proportion of sons that inherited the maternal X chromosomes from the total number of genotyped males ^20^, as only sons that have a maternal X are derived from the fertilization of a nullo-X sperm.

### Structural annotation of the genome

A comprehensive repeat library was produced for the genome assembly using multiple repeat-finding programs. The pipeline was based on the TransposableELMT wrapper script (https://github.com/PlantDr430/TransposableELMT). Repeats were identified using RepeatModeler v2.0.1 ^66^, TransposonPSI v08222010 ^67^, LTRfinder v1.0.7 ^68^ and LTRharvest ^69^(implemented in GenomeTools v1.6.1 ^70^. To limit the identification of false positives, the LTRharvest output was post-processed with LTRdigest ^71^ (implemented in GenomeTools v1.6.1 ^70^). The resulting libraries were combined, classified using RepeatClassifier v2.0.1 (part of the RepeatModeler v2.0.1 package ^68^ and redundancy removed using USEARCH ^72^ based on 90% similarity. The non-redundant custom library was used to soft mask repeat regions in the assembly with RepeatMasker v4.1.0 ^73^.

Gene predictions were made using the *ab initio* and evidence-driven gene predictors GeneMark-ES ^74^, SNAP (implemented in Maker2) ^75^, Maker2 ^76^ and Augustus^77^(trained with BUSCO v5.1.3 ^78^).

An *A. freiburgense* transcriptome assembled using Trinity ^79^, the *C. elegan*s protein database (UP000001940_6239), and the UniProt/Swiss-prot database (uniprot_sprot.fasta) were used as evidence-based inputs for the first round of Maker2. The resulting output was then used to train Augustus and SNAP. The unmasked genome was used as input into GeneMark-ES. Finally, the outputs from the first round of Maker2, Augustus, SNAP, and GeneMark were used as inputs in Maker2. The gene predictions from the second round of Maker2 were used for our analyses. The annotation was later lifted over to the final chromosomal-scale assembly using ’flo’ using chain files ^80^.

### Functional annotation of the genome

The functional annotation of the genome was conducted using the Blast2Go ^81^. A local Blast search of the *A. freiburgense* predicted proteins was conducted using a searchable database of the Swissprot proteins (swissprot.gz downloaded from NCBI July 2021) prepared within the Blast2Go software. The same predicted proteins were also analyzed with InterProScan (within the Blast2Go software) and the annotations merged to give the final functional annotation.

The completeness and quality of the predicted protein-coding genes were assessed using BUSCO on the transcript dataset in genome mode with gene discovery via Augustus against the nematode database (nematoda_odb10) (Table S9).

### Mitochondrial genome

The mitochondrial scaffold was identified by BLASTn (Version 2.9.0+)^53^, using the *A. rhodense* mitochondrial genome as a reference database. Only one scaffold was identified as the mitochondrion. This scaffold was analyzed and annotated using MITOS ^82^, which revealed a large duplication, probably due to a misassembly due to the mitochondrial genome being circular. We used the pair-wise local aligner Water ^83^ to identify precisely the junctions of the duplicated region and removed it using the subseq function of seqkit (version 0.16.1) ^84^. The curated mitochondrial genome was re-integrated into the assembly and replaced with the un-curated one. The annotations associated with the un-curated mitochondrial genome were removed and replaced by the MITOS annotations.

### Pooling DNA from lines with similar phenotypes into discrete pools and sequencing

Equal amounts of DNA from 10 RIAIL lines with the same phenotype were mixed to create pools of DNA from High Male lines (HM-Pools) and Low Male lines (LM- Pools). Two pools for each category were prepared, each containing 1.5 µg of DNA. The RIAIL lines used for the first and second HM-Pools were 23, 24, 26, 28, 29, 30, 33, 45, 57, 61 and 35, 42, 46, 55, 56, 59, 63, 65, 72 and 97, respectively. The RIAIL lines used for the first and second LM-Pools were 2, 5, 9, 13, 14, 17, 20, 54, 74, and 77, and 6, 8, 15, 18, 25, 27, 37, 40, 69, and 95, respectively. The quality of DNA was examined by running 1 μl of DNA on 1.8 % agarose gel, and concentration was measured using a Qubit fluorometer. Five DNA samples; 2 HM-pools, 2 LM-pools, and DNA from APS14 maternal strain, were sequenced. Sequencing libraries were generated using TruSeq DNA nano gel free at the GenePool facility at the University of Edinburgh on an Illumina HiSeq platform to generate 150-bp paired-end reads, with an insert size estimation of 350 bp.

### Variants calling for APS14 and next-generation sequencing Bulk segregant analysis (NGS-BSA)

The quality of the reads was assessed using FastQC software to have an overview of the reads’ quality ^85^. Paired-end reads were cleaned using Skewer using the following parameters"-n -Q 20 -l 51 -t 32 -m pe" ^86^. The quality of reads was reexamined after cleaning using FastQC software. Then, BWA program was used to align short reads from each pool and APS14 strain separately to the APS7 reference genome ^55^. Alignment files in SAM format were converted to BAM files using the samtools “view” command ^56^. BAM files from the HM-pools and the LM-pools were merged to produce a single BAM for each phenotype and were subsequently sorted using samtools ^56^. Variants from sorted BAM files of both pool samples and APS14 strain were called Individually using “Haplotypecaller" from GATK (Genomic Analysis Toolkit) ^25^. Low-quality variants were filtered out with ’VariantFiltration’ provided by GATK tools using default parameters ^25, 57–59^. Variants were filtered further using bcftools based on Genotype Quality >= 30 and depth >= 10 while keeping homozygous sites only ^60^. SnpEff (v5.0.1) ^26^ and SnpSift (v4.3.1) ^87^ were used to annotate and predict the effects of the genetic variants between APS14 and APS7 reference genome. Large structural variants between APS14 and APS7 genomes were identified using the Parliament2 pipeline ^88^. Structural variants were called using the following software: Breakdancer ^89^, CNVnator, ^90^, Manta ^91^, Lumpy ^92^ and DELLY ^93^. To avoid false positive calls we only considered structural variants that were called with more than one software.

Pools’ variant calling files were merged into one file using GATK "GenotypeGVCFs” command ^25^. Joint genotyping using “GenotypeGVCFs" combined all SNPs and indels records from both pools to produce correct genotype likelihood outputting a single combined variant calling file (VCF) ^94^. Variants were filtered using vcftools ^60^ removing all sites with missing values, removing all Indels, and keeping only biallelic sites. In total 334,230 SNPs were kept. The joint variant calling file was converted to a table using "VariantToTable" command provided by the GATK ^25^. Regions displaying differences between the HM-pools and LM-pools were identified using the R-package "QTLseqr” ^95^. QTLseq analysis for NGS-BSA was used to calculate the allele frequency difference (SNP-index) from the allele depth at individual SNP ^24^.

Candidate regions were identified by setting a sliding window size to 1 Mb, and the number of SNPs was counted in that window. Within each sliding window, a tricube-smoothed delta-SNP-index was calculated by constant local regression. A simulation was performed where the delta-SNP-index per bulk was calculated and simulated over 1000 replications based on RILs F2 population with a bulk size of 20.

Confidence intervals at 95% and 99% were estimated using the quantile from the simulation. An alternative approach, G-statistics, was used to identify significant QTLs from BSA ^96^. G-statistics was calculated genome-wide, and a tricube-smoothed G-statistics (G’) was predicted in a sliding window of 1M bp. P-values were estimated and adjusted (Benjamini-Hochberg method), and negative log10- were calculated from (G’). Significant regions were identified using a genome-wide false discovery rate of 0.1 ^97^.

### QTL analysis

The QTL analysis was conducted using rqtl on all the markers from all the scaffolds before they were organized into linkage groups to include markers with high segregation distortion that were omitted from the genetic linkage map and to include all the scaffolds, including those that were excluded from the final chromosomal assembly ^61, 98^. The 264,068 markers dataset imported into rqtl as RILs (see section "Construction of high-density genetic linkage map") were filtered using the rqtl function ‘drop.markers’ by removing markers that are not present in 90% of the data allowing only 10% of missing values and redundancy in the data set was reduced by removing markers that shared similar genotypes. All the markers with high segregation distortion were retained in the final data set of 18,513 markers from the 100 samples on 28 scaffolds. Markers segregation distortion is a natural phenomenon and the inclusion of distorted markers in QTL mapping increases the power of detecting a QTL ^98, 99^. The interval mapping genotype probability was calculated using the kosambi map function with a 1 CM step size and an error probability of 0.001. A whole-genome scan on all RIAILs was performed with a single QTL model using Haley–Knott regression, with the percentage of male progeny after an outcross for each RIAIL as a phenotype ^100^. The result from the first scan was permuted 1000 times to obtain a genome-wide LOD score (logarithm of odds) significance threshold ^101^. Genome-wide thresholds were obtained at p values < 0.05 and < 0.2 corresponding to a LOD score of 2.98 and 2.35 respectively. Interval estimates of the location of the QTL were obtained via 1.5-LOD support intervals, which could be calculated via the function (lodint). Three QTLs on three different scaffolds belonging to the X chromosome were detected from the single scan.

Scanning for the likelihood of interacting QTLs using a two-dimensional genome scan, for all pairwise combinations of intervals, was impossible due to a large number of markers ^102^. Hence, a multiple QTL analysis using QTLs identified from the single scan was performed. Briefly, a qtl object for multiple qtl analysis was created containing all the identified QTLs from the single scan using the rqtl function (makeqtl). QTLs were fitted into an additive model using the ’addinit’ function, which adds one interaction at a time in the context of a multiple interval mapping (MIM) model, model formula: y ∼ QTL1 + QTL2 + QTL3. An interaction was detected between the three QTLs identified from the single scan at p-value (Chi2) < 0.05. The interaction was expected as the three QTLs were on scaffolds assembled adjacent to each other on the X Chromosome. The estimate of the qtl peak location was refined using the ’refinqtl’ function. Finally, the function ’addqtl’ was used to scan for an additional QTL to be added to the model. Markers outlining the start and the end of QTL regions identified on scaffolds were overlapped with the genetic linkage map to determine the physical location of the QTL region on the final genome assembly. The three QTLs were adjacent to each other on the X linkage group, therefore they were merged into a single QTL. In a separate analysis, the 18,513 markers dataset from 22 scaffolds (nonredundant and includes all markers with segregation distortion) used to scan for a QTL was merged with the genetic linkage map to obtain the genetic position of each marker ^20^. The analysis was repeated using all the steps above. A single QTL was detected on the X chromosome confirming the three identified QTLs in the earlier analysis are part of one continuous X chromosome QTL.

## Availability of the data

We deposited all sequencing data at the European Nucleotide Archive (ENA) under BioProject numbers PRJEB55706 (Illumina genomic Data), PRJEB60474 and PRJEB50372 (Illumina transcriptomic data), and PRJNA640723 (PacBio data).

Furthermore, we uploaded the genome assembly, annotation, and predicted protein dataset to NCBI with accession PRJNA947217.

