## Supplemental Figures and Tables for "The contribution of an X chromosome QTL to non-Mendelian inheritance and unequal chromosomal segregation in *A. freiburgense*"

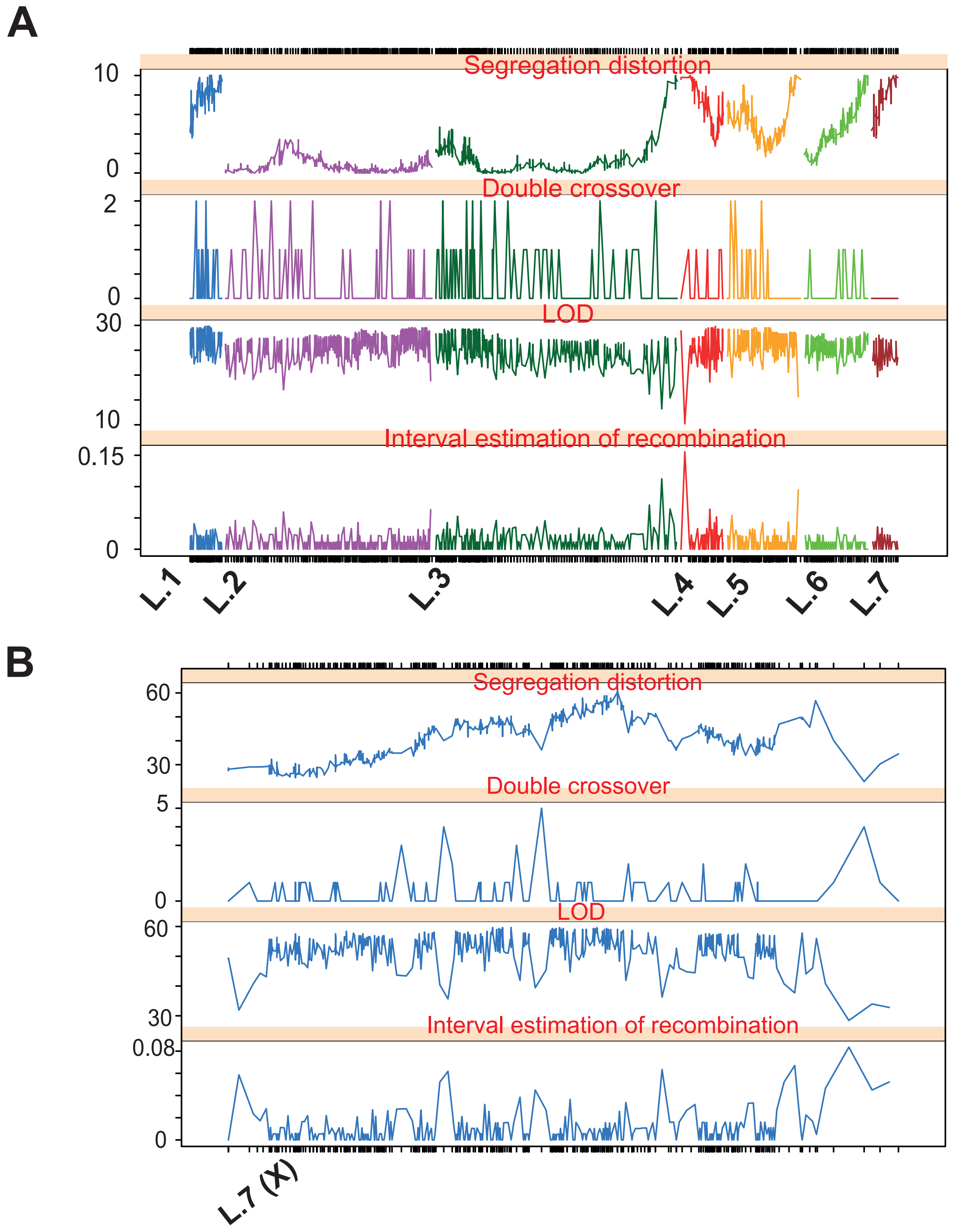


**Figure S1. Comparison of the metrics of individual markers in the initial genetic linkage map (GLM) and the independent GLM constructed for the X chromo-some.** The initial GLM (**A**), constructed using only homozygous markers, did not detect any double crossovers on L.7 (X chromosome). In contrast, the inde-pendent GLM (**B**), which was constructed using both homozygous and hete-rozygous markers, detected double crossovers and resulted in an increase in the size of the X chromosome. The interval estimation of recombination and marker segregation distortion is generally low in both GLMs. Ticks on the x-axis represent markers, and the y-axis shows the metrics for each plot title in differ-ent panels. Plots were produced using the ASMap R package after the con-struction of the genetic linkage map.

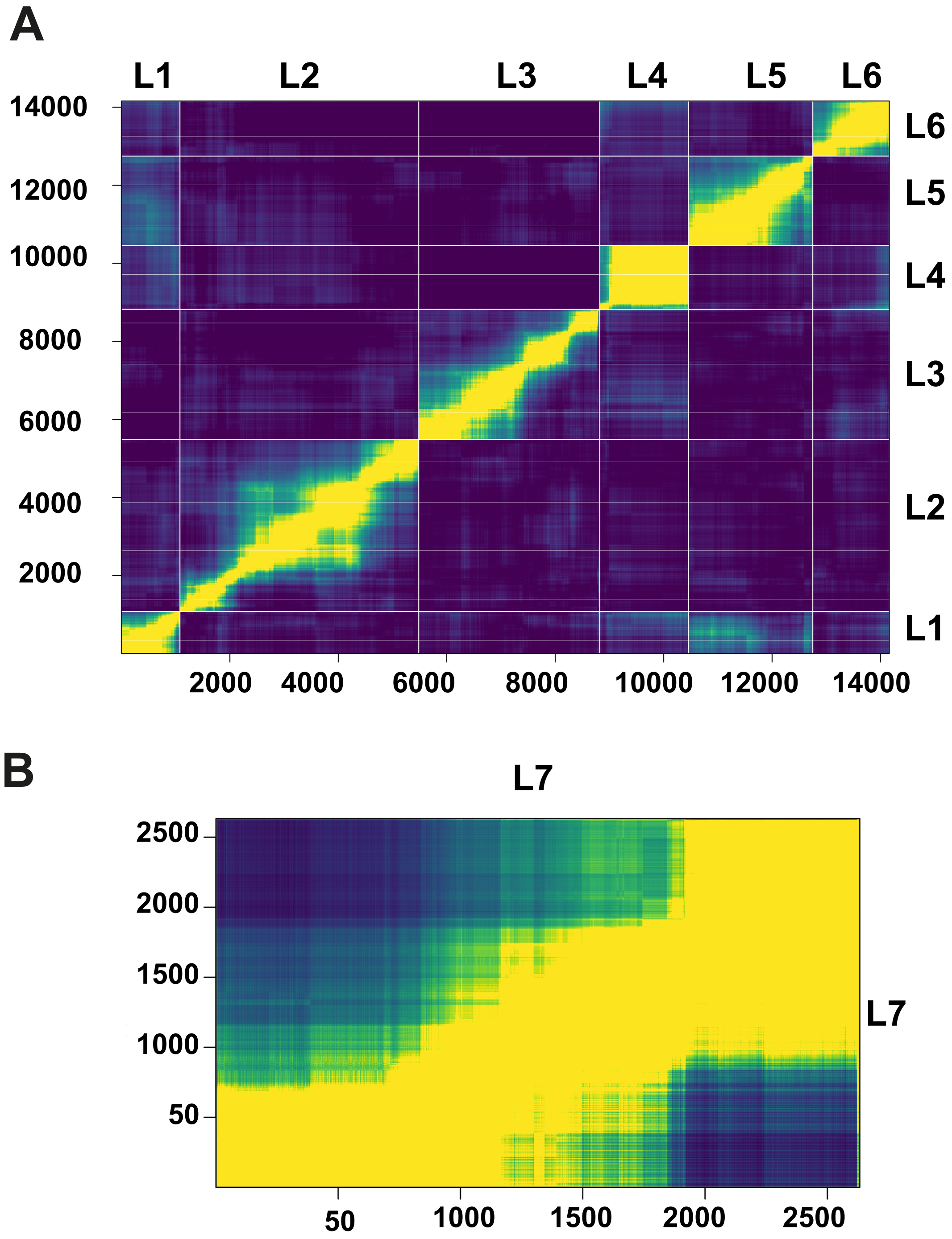


**Figure S2.** Heatmaps for autosomal and sex chromosome genetic linkage maps. The heatmaps show the clear clustering of markers for each linkage group. The autosomal heatmap (**A**) was constructed using only homozygous markers, whereas the heat map for the X chromosome L.7 (**B**) was built separately, using homozygous and heterozygous markers.

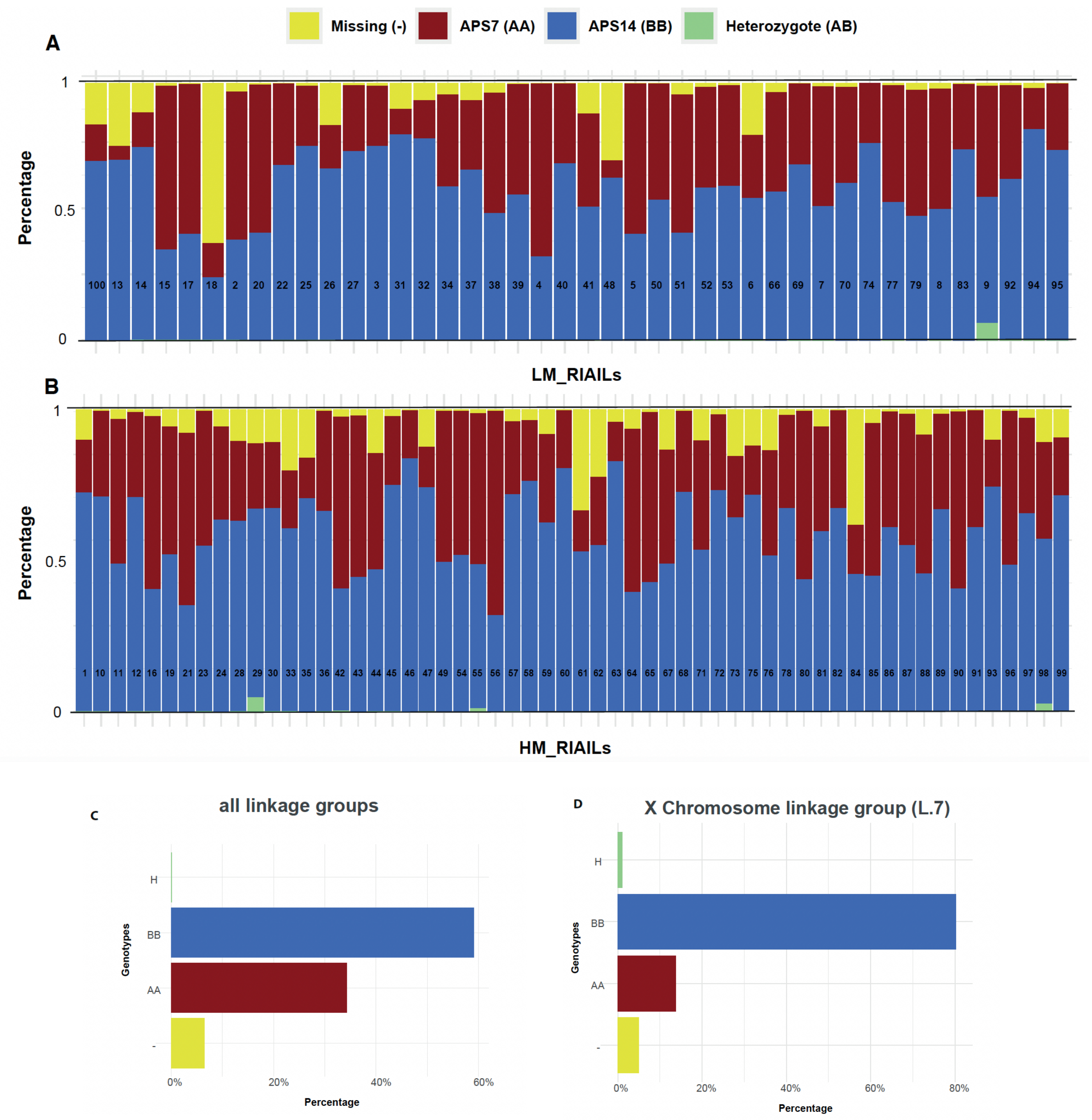


**Figure S3. Individual RIAILs Genotype ratio (A and B), percentage of genotypes in the genetic linkage map (C) and the X chromosome (D).** The genotype composition of individual RIAILs in the X-axis separated by phenotypes LM-RIAILs (A) and HM-RIAILs (B), was plotted as a ratio on the y-axis. The ratio of missing markers (yellow) in each individual is generally low. There is no particular pattern of genotype ratio inheritance associated with RIAILs phenotype. The percentage of genotypes in the overall genetic linkage map from all lines indicates that lines inherited more markers from APS14 parent (BB) 59% than the APS7 parent (AA) 34%, and the percentage of missing markers is only 6% (C). The majority of markers on the X chromosome (L.7) in all lines are inherited from APS14 (BB) 80% and the percentage of heterozygous markers is 1.1% (D).

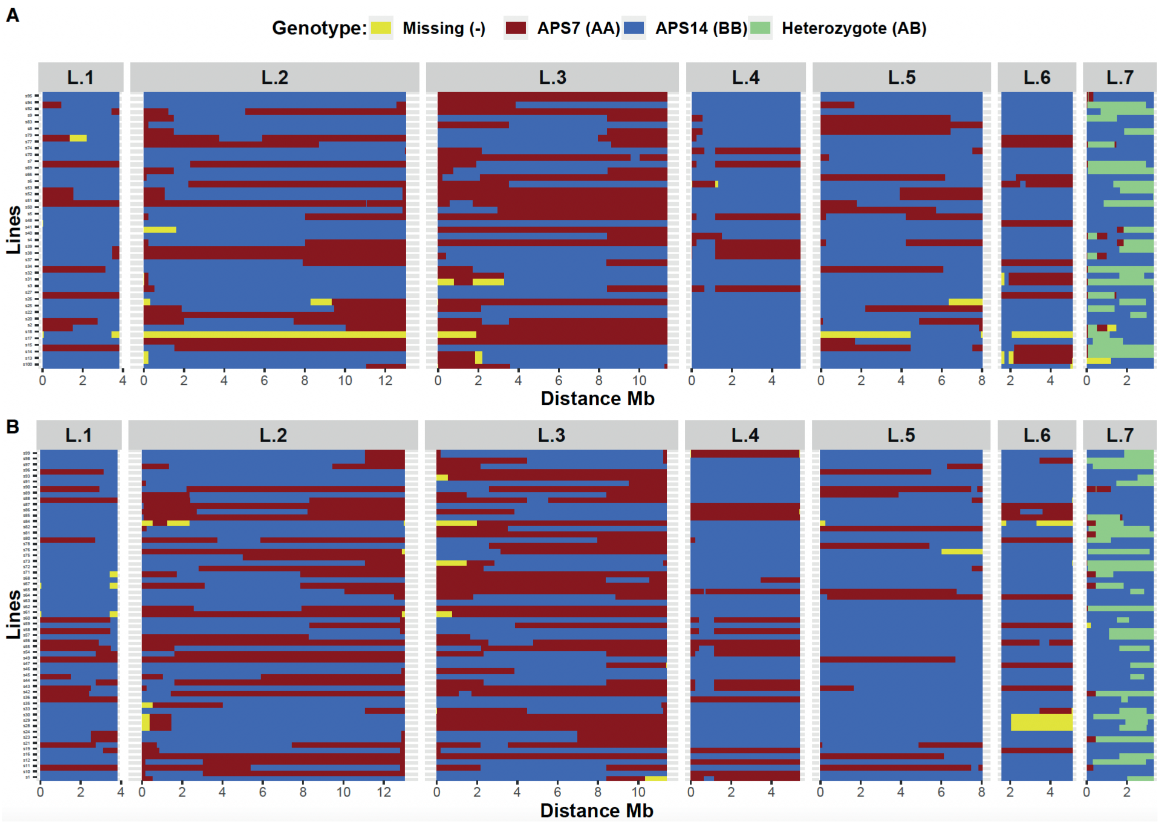


**Figure S4. Individual RIAILs parental background separated by phenotype.** Genotypes were plotted across the 7 linkage groups, each linkage group is in a separate panel on the (x-axis), genome-wide recombination in each line is represented by the mosaic blocks of colors. RIAILs (y-axis) parental background, determined from markers used to construct the genetic linkage map were plotted across the genetic distance in Mb separated by phenotype (A) HM-RIAILs (B) LM-RIAILs.

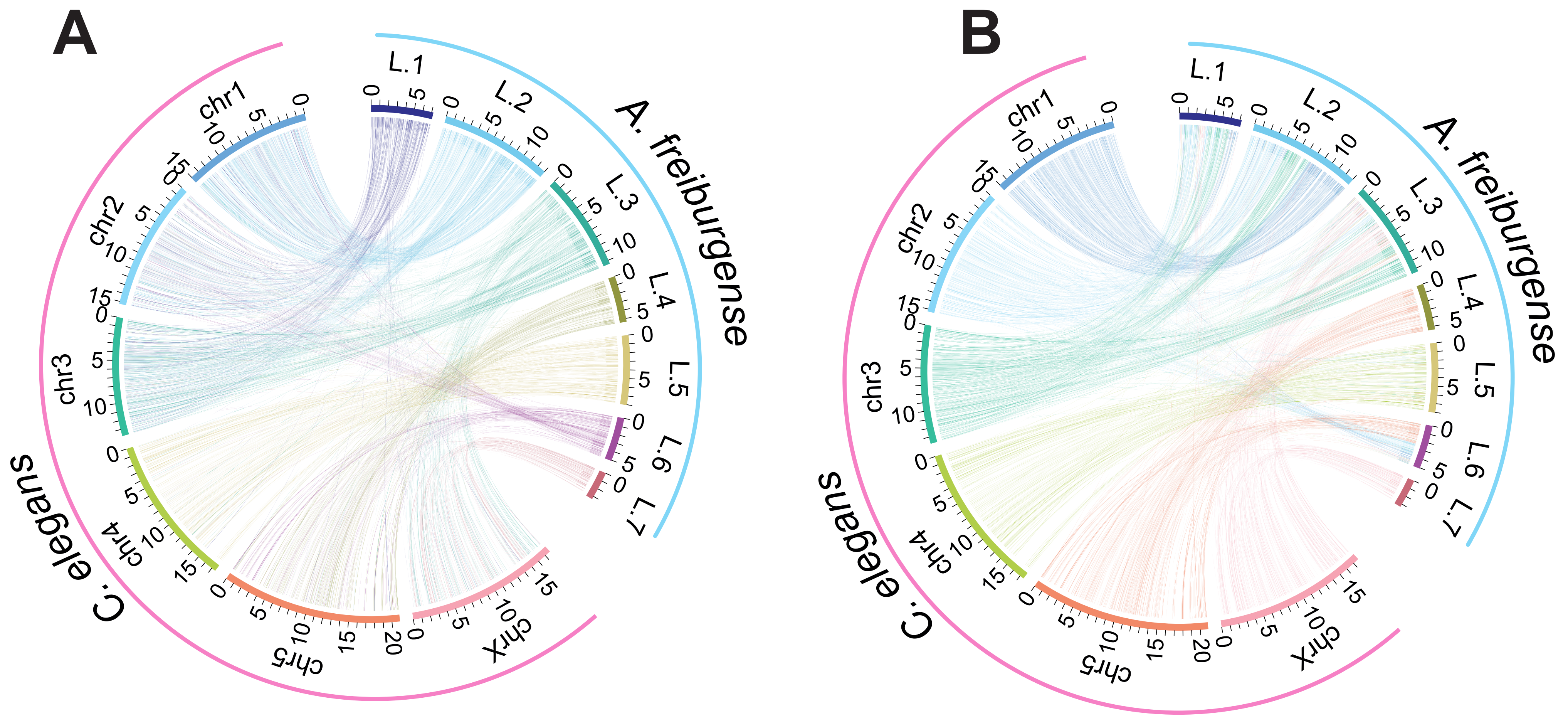


**Figure S5. Macrosynteny relationship between *A. freiburgense* and *C. elegans*.** (A) Color mapping according to *A. freiburgense* chromosomal-scale linkage groups. (B) Color mapping according to *C. elegans* chromosomes. Each line links an orthologous pair of genes.

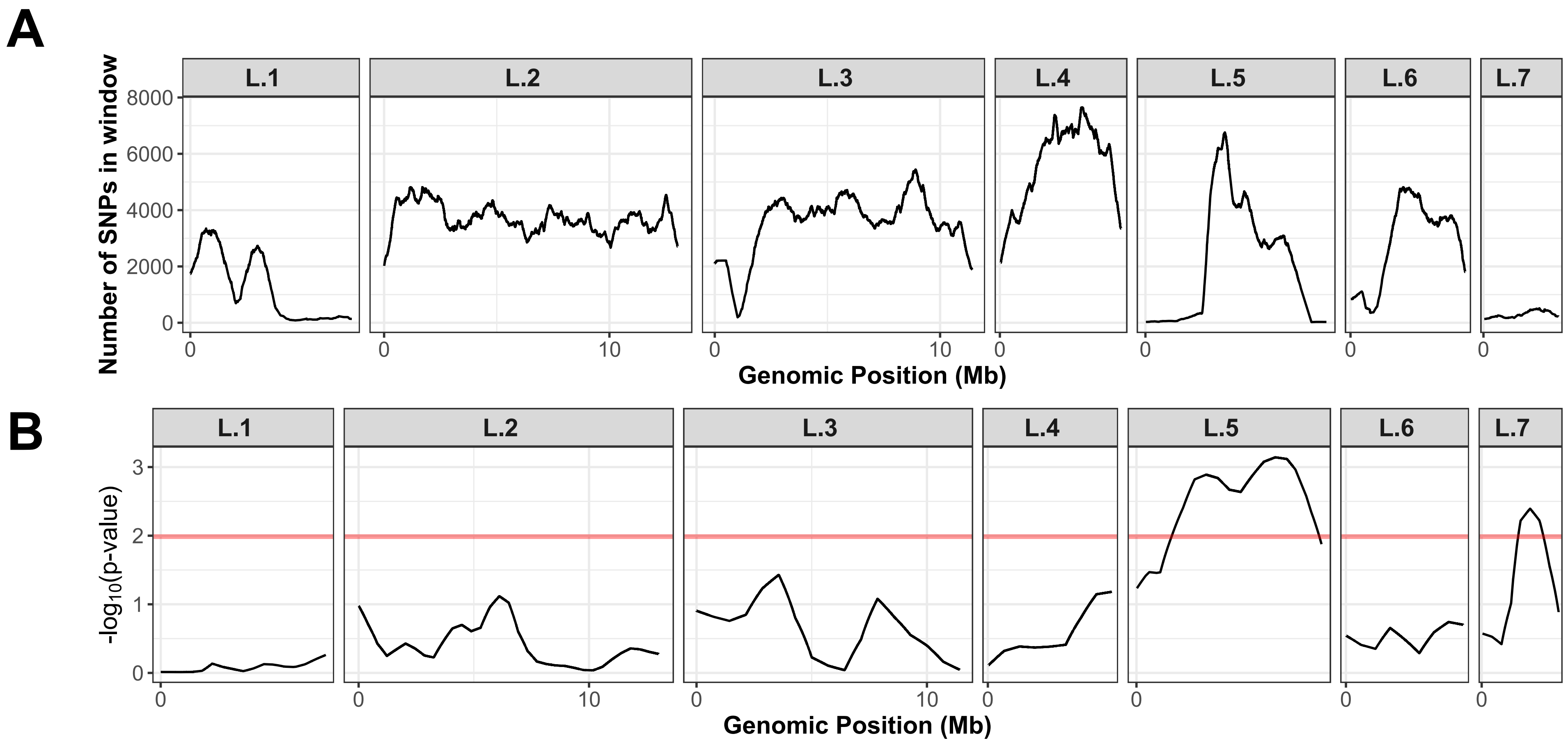


**Figure S6. Bulk segregant analysis (BSA) identified two candidate regions using tricube-smoothed G statistic (G') .** For the 334230 SNPs detected from both pools, NGS-BSA calculated SNP number **(A)** in a sliding window of 1M bp and tricube-smoothed G statistic (G') in a sliding window of 1M bp **(B)**. Contiguous regions with a q-value above 0.1 were identified as regions of significance.

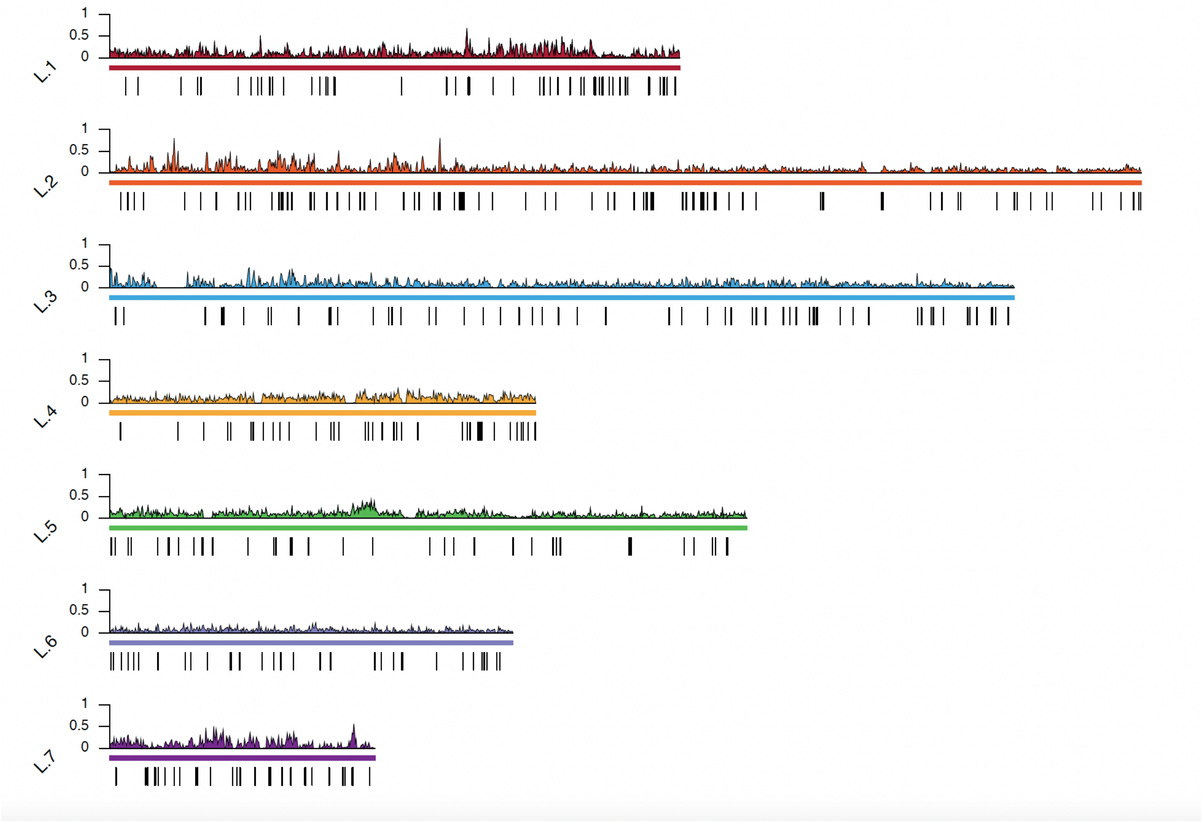


**Figure S7. APS14 Indels density and structural deletions.** Indels density was plotted in a density plot color coded for each chromosome using a window size of 10000 bp. APS14 structural deletions were plotted in black bars representing the gaps in APS14 genome.

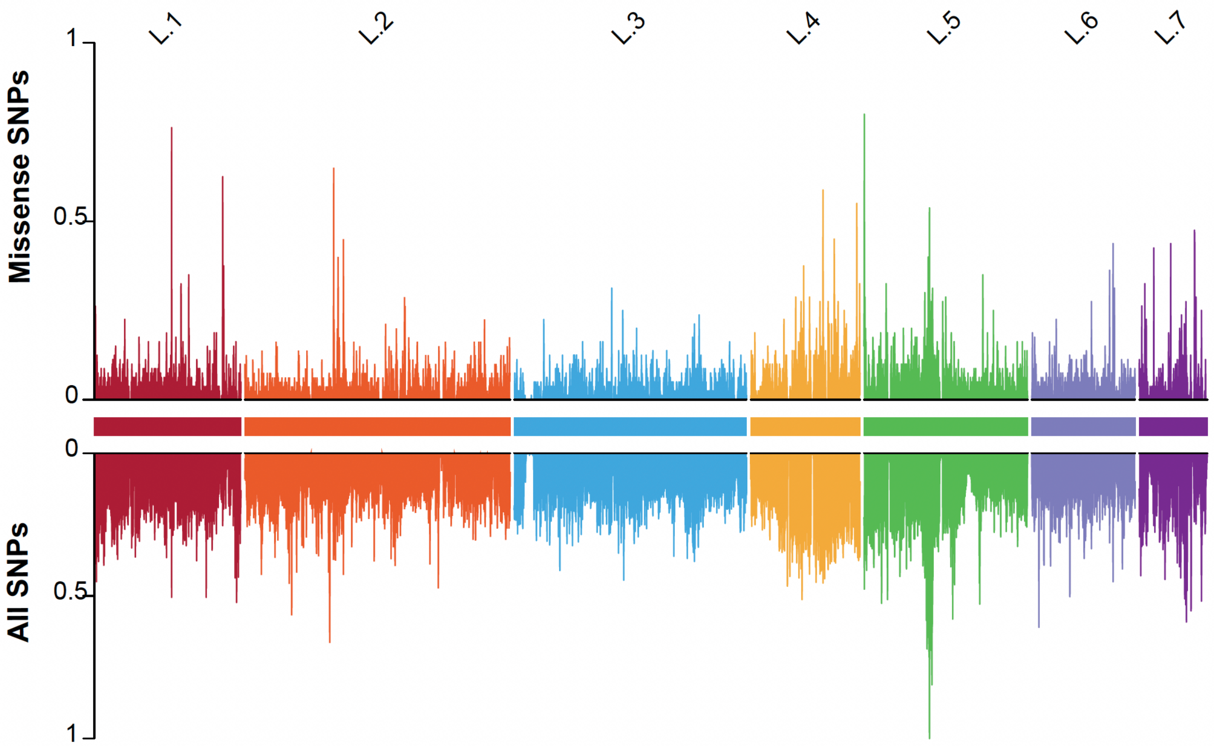


**Figure S8. APS14 SNPs density.** Density of SNPs on a window size of 10,000 bp represented in ratio from 0 to 1 plotted for each chromosome color coded.

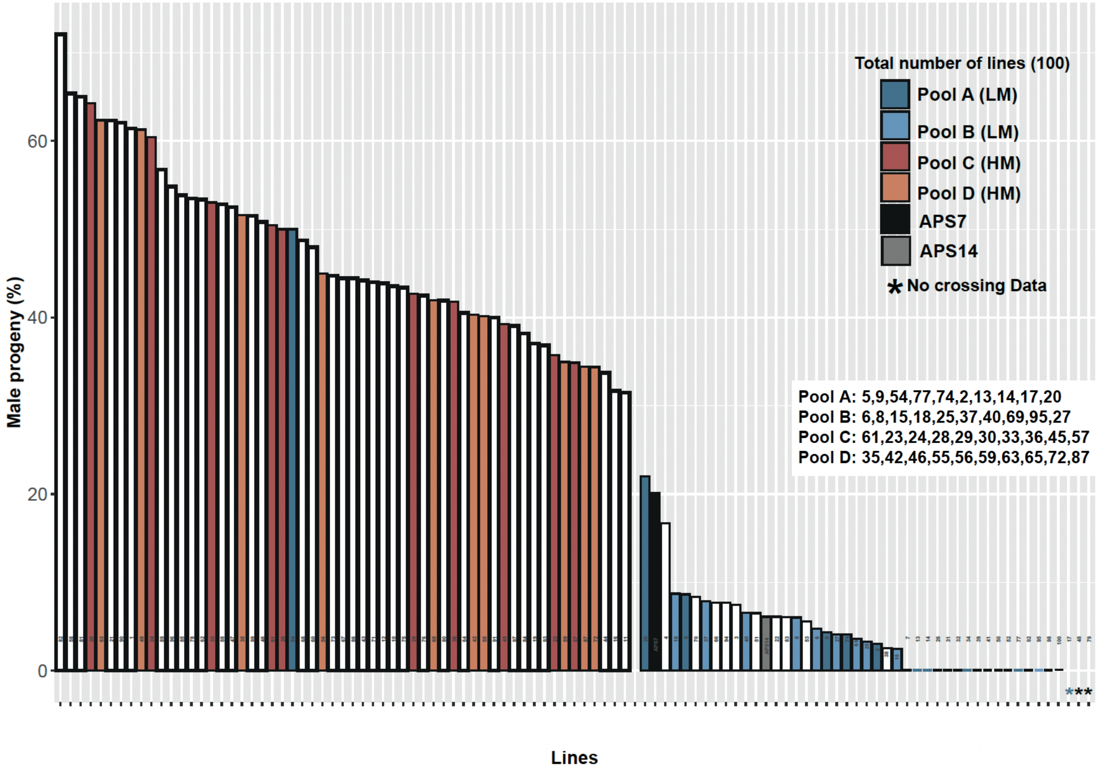


**Figure S10. Lines used in the bulk segregant analysis (BSA).** The percentage of males produced when a male from each line is crossed with APS7 mother is represented by a bar chart. DNA from the lines highlighted were pooled to constitute 4 pools; 2 LM pools (A and B) and two HM pools (C and D). Line 54 was mistakenly pooled with LM-pool despite it being a HM line. This mistake would have not biased the BSA analysis since most of the samples in the pool are still LM lines.

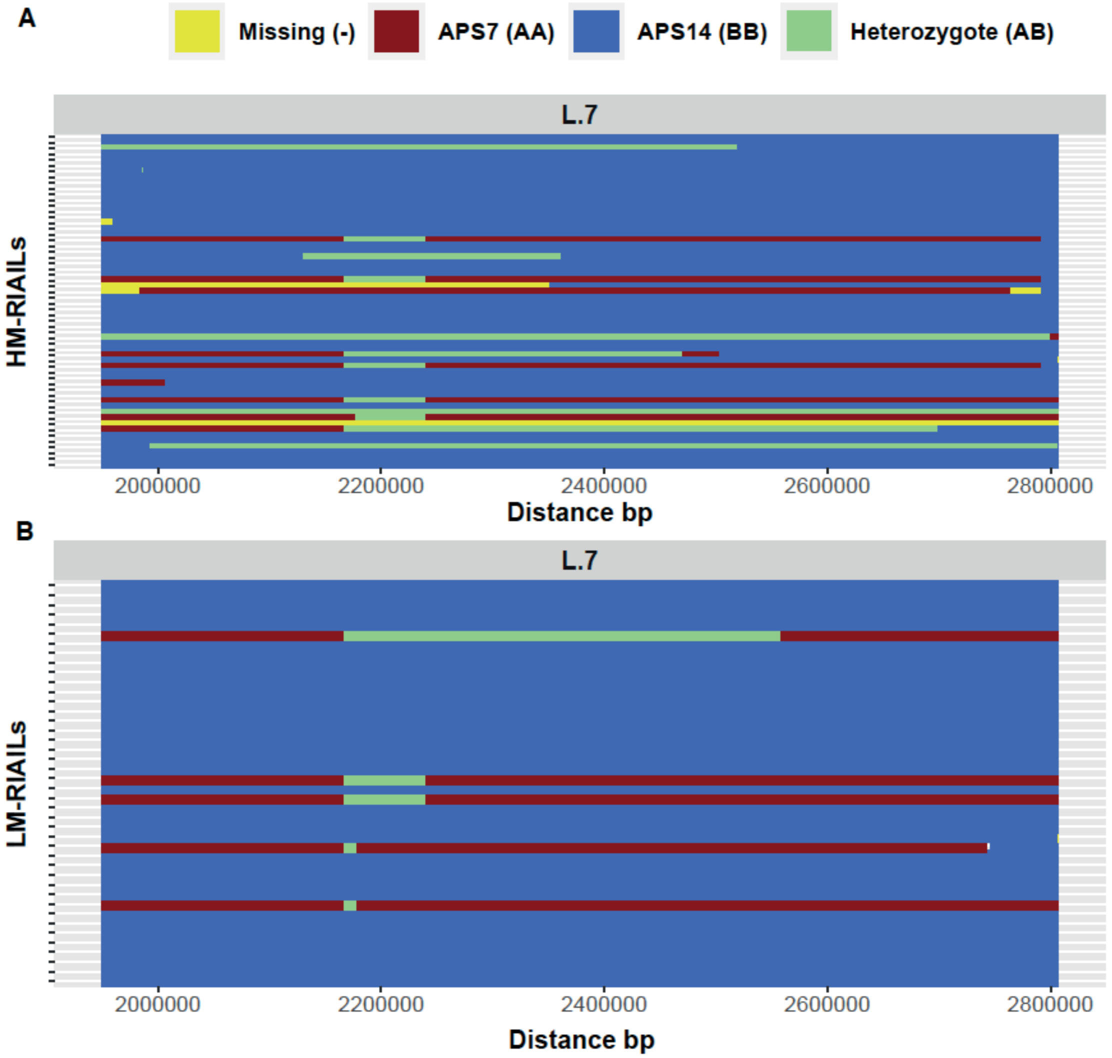


**Figure S9. Parental background of genotypes within the QTLregion.** There is no apparent pattern of genotype inheritance associated with RIAILs phenotype.

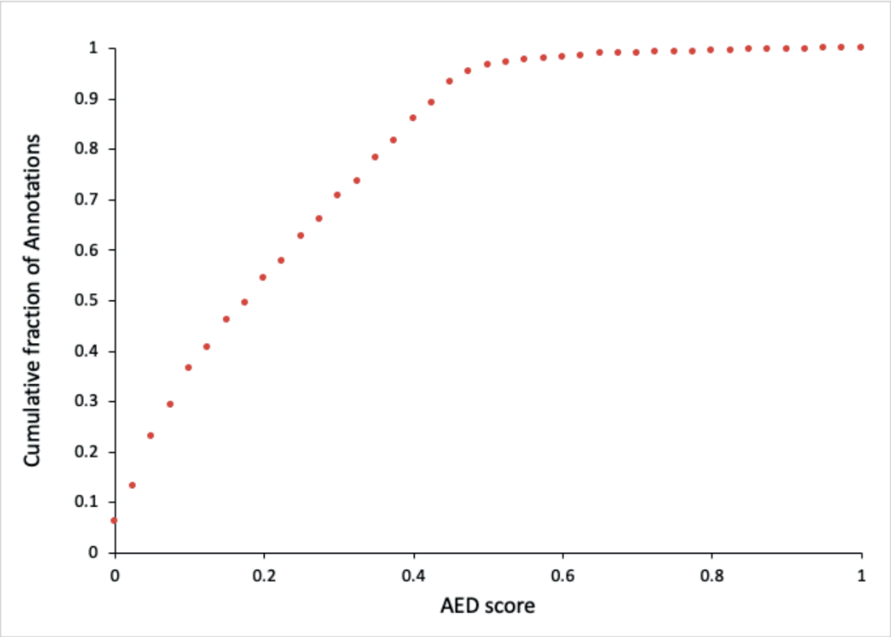


**Figure S11. Cumulative Annotation Adit Aistance (AED) of the *A. freiburgense* genome annotation.** Annotation edit distance (AED) is a general measure of how well the predicted gene is supported by external evidence (UniProt protein and mRNA sequences) and was conducted using the AED_cdf_generator.pl script (ttps://github.com/mscampbell/Genome_annotation/blob/master/AED_cdf_generator.pl) using Genome_Annotation_v2.maker_genes_only.gff as input.

**Table S1. Comparison of the X chromosome assembly when only homozygous markers versus when heterozygous markers are also included.** Including heterozygosity when linking X chromosome scaffolds increases X chromosome contiguity. Percentages reported excludes the percentage of missing markers in each linkage. Percentages of genotypes including the missing markers for the X linkage included in the final genome assembly are highlighted in Figure S3.

| X chromosome | Number of markers | Assembly length in bp | %AA | %BB | %AB |
|---|---|---|---|---|---|
| Assembly with only homozygous markers | 724 | 1,587,724 | 23.2 | 76.8 | Not included |
| Assembly by including heterozygous markers | 2,632 | 3,358,134 | 14.5 | 84.4 | 1.2 |

**Table S2. Summary of the physical genome assembly of *A. freiburgense***

| Summary of genetic linkage map | | | |
|---|---|---|---|
| Linkage Groups | 7 | | |
| Markers (unique) | 16,792 | | |
| Markers per Mb | 313.3 | | |
| N50 Scaffolds | 7 | | |
| Scaffolds | 29 | | |
| Scaffolds with 3 markers | 1 | | |
| Scaffolds with >=4 markers | 28 | | |
| Total bases | 53,604,205 (97.0%) | | |
| Summary for consensus map after genetic linkage map anchored, ordered and oriented | | | |
| | Anchored | Oriented | Unplaced |
| Markers (unique) | 16,787 | 16,324 | 5 |
| Markers per Mb | 313.3 | 308.9 | 3.0 |
| N50 Scaffolds | 7 | 7 | 7 |
| Scaffolds | 28 | 24 | 47 |
| Total bases | 53,572,943 (96.9%) | 52,848,025 (95.6%) | 1,689,119 (3.1%) |

**Table S3. Contiguity and BUSCO score of the *A. freiburgense* genome assembly.**

| Assembly stage | | Pacbio assembly (version. 1) | Genetic linkage map, final genome assembly |
|---|---|---|---|
| Genome Contiguity metrics | contigs | 75 | 7 and 1 unplaced contig |
| | Total length Mb | 55.3 | 55.3 |
| | Largest contig | 13,037,505 bp | 7,206,650 bp |
| | N50 (n = contigs >= N50) | 3,036,117 (n = 7) | 8,053,364 (n = 3) |
| | N70 (n= contigs >= N70) | 2,305,519 (n = 11) | 7,206,650 (n = 4) |
| | N90 (n= contigs >= N90) | 669,147 (n = 18) | 5,099,687 (n = 6) |
| BUSCO score* | C [S] | 2791 (89.2%) [2757 (88.1%)] | |
| | D | 34 (1.1%) | |
| | F | 46 (1.5%) | |
| | M | 294 (9.3%) | |

*C: Complete BUSCOS (include single-copy and duplicated BUSCOS), D:Complete and duplicated BUSCOs, F: Fragmented BUSCOs, M: Missing BUSCOs. Busco v5.2.2 was used on the nematoda-odb10 database (3131 BUSCO groups)

**Table S4. Classification of repeat elements in the *A. freiburgense* genome using RepeatMasker.** A total of 18.2% of the genome (10,058,403 bp) was identified as repeat elements and was masked for annotation purposes.

| | Number of Elements | Length occupied (bp) | Percentage of total sequence (%) |
|---|---|---|---|
| **Retroelements** | 862 | 2066244 | 3.74 |
| SINES: | 45 | 39433 | 0.07 |
| LINES: | 51 | 122612 | 0.22 |
| L2/CR1/Rex | 11 | 10423 | 0.02 |
| R2/R4/NeSL | 24 | 51174 | 0.09 |
| L1/CIN4 | 16 | 61015 | 0.11 |
| LTR elements: | 766 | 1904199 | 3.45 |
| BEL/Pao | 277 | 812726 | 1.47 |
| Gypsy/DIRS1 | 467 | 969111 | 1.75 |
| **DNA transposons** | 569 | 373476 | 0.68 |
| hobo-Activator | 149 | 87123 | 0.16 |
| Tc1-IS630-Pogo | 60 | 30344 | 0.05 |
| PiggyBac | 1 | 645 | 0.00 |
| **Rolling-circles** | 106 | 85632 | 0.15 |
| **Unclassified** | 3162 | 3977979 | 7.20 |
| **Total interspersed repeats** | | 6417699 | 11.61 |
| **Small RNA** | 1003 | 1939812 | 3.51 |
| **Satellites** | 162 | 499587 | 0.90 |
| **Simple repeats** | 18218 | 839394 | 1.52 |
| **Low complexity** | 5099 | 276279 | 0.50 |

**Table S5. Distribution of Nigon elements in *A. freiburgense's* linkage groups.** Percentages of each Nigon Element in each chromosome. Percentages above 20% were highlighted in bold.

| | Nigon A | Nigon B | Nigon C | Nigon D | NigonE | NigonN | Nigon X |
|---|---|---|---|---|---|---|---|
| L.1 | 13.5 | **29.6** | **26.3** | 0 | 1.1 | 11.1 | 1.1 |
| L.2 | **80.3** | **35.5** | **31.2** | 0 | 0 | 0 | 1.1 |
| L.3 | 0 | 9.4 | **42.5** | 0.8 | 0 | **83.4** | 1.1 |
| L.4 | 0.2 | 0 | 0 | 0 | **63.5** | 0 | 0 |
| L.5 | 0.4 | 0 | 0 | **98.7** | 0 | 0 | 2.2 |
| L.6 | 5.6 | **25.5** | 0 | 0.4 | **35.4** | 5.5 | 1.1 |
| L.7 | 0 | 0 | 0 | 0 | 0 | 0 | **93.4** |
| percentage | 100 | 100 | 100 | 100 | 100 | 100 | 100 |

**Table S6. The Bulk Segregant Analysis (BSA) identified two candidate regions on L.5 and L.7.** NGS-BSA calculated tricube-smoothed delta-SNP-index and tricube-smoothed G statistic (G') in a sliding window of 1M bp.

| LG | qtl | Start | End | Length | Number of SNPs in window | Average number of SNPs in a Mb | Peak Delta SNP | Position of Peak Delta SNP |
|---|---|---|---|---|---|---|---|---|
| L.5 | 1 | 1,523,958 | 7,906,199 | 6,382,241 | 16560 | 2595 | 0.44 | 6,029,576 |
| L.7 | 2 | 1,606,040 | 2,683,704 | 1,077,664 | 494 | 458 | 0.33 | 2,104,550 |

| qtl | Avgerage Delta SNP | Maximum Gprime | Position of Max Gprime | Mean of Gprime | Sd of Gprime | AUCaT | Mean Pval | meanQval |
|---|---|---|---|---|---|---|---|---|
| 1 | 0.37 | 372.12 | 6,029,576 | 306.48 | 38.23 | 892,266,716 | 0.001 | 0.023 |
| 2 | 0.30 | 209.56 | 2,104,550 | 189.01 | 16.45 | 44,540,969.6 | 0.006 | 0.055 |

**Table S7. Number and percentage of APS14 variants by their type, functional class, effect and regions as predicted by SnpEff.** The percentage of each variant type is calculated per the total from each group.

| Number variants by type | | |
|---|---|---|
| **Type** | **Count** | **Percentage** |
| SNP | 298,505 | 84.1 |
| INS | 30,673 | 8.6 |
| DEL | 25,527 | 7.2 |
| Total | 354,705 | |

| Number of effects by functional class | | |
|---|---|---|
| **Type** | **Count** | **Percentage** |
| MISSENSE | 14,946 | 27.6% |
| NONSENSE | 202 | 0.4% |
| SILENT | 39,041 | 72.0% |
| Total | 54,186 | |
| Missense / Silent ratio: 0.3828 | | |

| Number of effects by type and region | | | | | |
|---|---|---|---|---|---|
| **Type** | | | **Region** | | |
| **Type (alphabetical order)** | **Count** | **Percentage** | **Type (alphabetical order)** | **Count** | **Percentage** |
| 3 prime UTR variant | 22,799 | 2.13% | DOWNSTREAM | 346,698 | 32.9% |
| 5 prime UTR premature start codon gain variant | 2,489 | 0.23% | EXON | 54,065 | 5.1% |
| 5 prime UTR variant | 19,146 | 1.79% | GENE | 4 | 0% |
| bidirectional gene fusion | 3 | 0% | INTERGENIC | 138,696 | 13.1% |
| conservative inframe deletion | 99 | 0.01% | INTRON | 107,751 | 10.2% |

| | | | | | |
|---|---|---|---|---|---|
| conservative inframe insertion | 105 | 0.01% | SPLICE SITE ACCEPTOR | 184 | 0.02% |
| disruptive inframe deletion | 166 | 0.02% | SPLICE SITE DONOR | 315 | 0.03% |
| disruptive inframe insertion | 175 | 0.02% | SPLICE SITE REGION | 17,620 | 1.6% |
| downstream gene variant | 346,702 | 32.36% | TRANSCRIPT | 80 | 0.01% |
| frameshift variant | 585 | 0.06% | UPSTREAM | 343,281 | 32.6% |
| gene fusion | 1 | 0% | UTR 3 PRIME | 22,781 | 2.1% |
| initiator codon variant | 5 | 0% | UTR 5 PRIME | 21,627 | 2.0% |
| intergenic region | 138,696 | 12.94% | Total | 1,053,102 | |
| intragenic variant | 10 | 0.00% | | | |
| intron variant | 123,992 | 11.57% | | | |
| missense variant | 14,887 | 1.39% | | | |
| non-coding transcript exon variant | 20 | 0.00% | | | |
| non-coding transcript variant | 69 | 0.01% | | | |
| splice acceptor variant | 188 | 0.02% | | | |
| splice donor variant | 325 | 0.03% | | | |
| splice region variant | 18,333 | 1.71% | | | |
| start lost | 45 | 0.00% | | | |
| start retained variant | 2 | 0% | | | |
| stop gained | 260 | 0.02% | | | |
| stop lost | 37 | 0.00% | | | |
| stop retained variant | 66 | 0.01% | | | |

| | | |
|---|---|---|
| synonymous variant | 38,982 | 3.6% |
| transcript ablation | 1 | 0% |
| upstream gene variant | 343,281 | 32.0% |
| Total | 1,071,469 | |

**Table S8. Structural deletions in the QTL region.** Large deletions were identified in the QTL region when structural variants were called using the Parliament2 pipeline.

| Chr | Start | End | Width |
|---|---:|---:|---:|
| chr7 | 2,010,139 | 2,012,502 | 2,364 |
| chr7 | 2,027,707 | 2,034,544 | 6,838 |
| chr7 | 2,172,524 | 2,175,910 | 3,387 |
| chr7 | 2,177,187 | 2,178,345 | 1,159 |
| chr7 | 2,179,699 | 2,185,972 | 6,274 |
| chr7 | 2,285,179 | 2,290,685 | 5,507 |
| chr7 | 2,465,641 | 2,467,931 | 2,291 |
| chr7 | 2,470,776 | 2,481,758 | 10,983 |
| chr7 | 2557771 | 2559052 | 1,282 |
| chr7 | 2,770,830 | 2,771,380 | 551 |
| chr7 | 2,772,220 | 2,776,919 | 4,700 |
| chr7 | 2,777,486 | 2,778,852 | 1,367 |

**Table S9. Raw data of *A. freiburgense* (strain APS7 and APS14) used in the study.**

| Accession | Type | Platform | Read metrics | Use | Number of reads |
|---|---|---|---|---|---|
| ENA Project PRJEB55706 | Genomic DNA | Illumina HiSeq | 150-bp paired-end reads, with an insert size estimation of 350 bp | Bulk segregant analysis | 1,002,297,576 |
| | Genomic DNA | Illumina HiSeq | 150-bp paired-end reads, with an insert size estimation of 350 bp | Genetic linkage mapping (Individual RIAIL) | 2,518,494,166 |
| ENA Project PRJEB61637 | Genomic DNA | Illumina HiSeq | 100bp pair-end library (insert size estimation of 250bp) | Genome correction and Polishing | 32,827,673 |
| | Genomic DNA | Illumina HiSeq | 100bp pair-end library (insert size estimation of 250bp) | Genome correction and Polishing | 39,391,272 |
| | Genomic DNA | Illumina HiSeq | 100bp pair-end library (insert size estimation of 250bp) | Genome correction and Polishing | 34,887,890 |
| | Genomic DNA | Illumina HiSeq | 125bp mate-pair library(insert size estimation of 3Kb) | Genome correction and Polishing | 97,192,878 |
| | Genomic DNA | Illumina HiSeq | 125bp mate-pair library(insert size estimation of 3Kb) | Genome correction and Polishing | 42,568,779 |
| | Genomic DNA | Illumina HiSeq | 125bp mate-pair library(insert size estimation of 5Kb) | Genome correction and Polishing | 80,016,124 |
| | Genomic DNA | Illumina HiSeq | 125bp mate-pair library(insert size estimation of 5Kb) | Genome correction and Polishing | 3,5417,072 |
| ENA Project PRJNA640723 | Genomic DNA | Pacbio | | Genome assembly | 1,652,896 |
| Experiment SRX8586548<br>Experiment SRX8586549 | Long RNA-seq | Pacbio | | Transcriptome Assembly | 15,415,711 |
| ENA Project PRJEB50372 | RNA-seq | Illumina HiSeq | 150 bp pair-end reads of female and hermaphrodite APS7 (inbred SB372) on day 2 of adulthood | Transcriptome Assembly | 292,110,832 |
| ENA Project PRJEB60474 | RNA-seq | Illumina HiSeq | 150 bp pair-end reads of female and | Transcriptome Assembly | 329,639,162 |

| | | | | | |
|---|---|---|---|---|---|
| | | | hermaphrodite JU1782 on day 2 of adulthood | | |
| | RNA-seq | Illumina HiSeq | 150 bp pair-end reads of SB372 hermaphrodites | Transcriptome Assembly | 47,840,008 |
| | RNA-seq | Illumina HiSeq | 150 bp PE reads of mixed stage SB372 | Transcriptome Assembly | 119,780,330 |

### Table S10. List of software and parameters

| Software | Version | Source | Parameter used | Reference |
|---|---|---|---|---|
| Allmaps | v1.0.9+4.g417e8d1b | https://github.com/tanghaibao/jcvi/wiki/ALLMAPS | 'merge' to produce .bed file<br>'Path' for scaffold ordering | [1] |
| ASMap | 1.0.4 | https://github.com/cran/ASMap | | [1] |
| bcftools | 1.9 | https://github.com/samtools/bcftools | | [2] |
| BLAST+ | 2.7.1 & 2.9.0 | https://blast.ncbi.nlm.nih.gov/Blast.cgi?PAGE_TYPE=BlastDocs&DOC_TYPE=Download | Blastn | [3] |
| Blast2Go Basic | 6.0.1 | https://www.biobam.com/download-blast2go/ | | |
| BUSCO (includes Augustus) | 5.2.2 | https://busco.ezlab.org/ | --augustus --long -m genome<br>-l nematoda_odb10 | [4] |
| BWA | 0.7.12-r1039 | http://bio-bwa.sourceforge.net/ | | [5] |
| canu | 1.6 | https://canu.readthedocs.io/en/latest/quick-start.html | genomeSize=55m<br>correctedErrorRate=0.030<br>minReadLength=3kb | [6] |
| Circos | | http://circos.ca/ | | [7] |
| FastQC | v0.11.5 | http://www.bioinformatics.babraham.ac.uk/projects/fastqc/ | | [8] |
| flo | | https://github.com/wurmlab/flo | | [9] |
| GATK | 3.8-0-ge9d806836 | https://gatk.broadinstitute.org/hc/en-us | | [10] |
| GeneMark-ES | 4.65 | http://topaz.gatech.edu/GeneMark/license_download.cgi | | [11] |

| | | | | |
|---|---|---|---|---|
| GenomeTools (includes LTRharvest and LTRdigest) | 1.6.1 | https://github.com/genometools/genometools | For LTRharvest<br>*Parameters to make the index with suffixerator*<br>-tis -suf -lcp -des -ssp -sds -dna -lossless<br>*Parameters to run LTRharvest*<br>-seqids yes<br>-md5<br>-tabout no | [12] |
| LTRfinder | 1.0.7 | https://github.com/xzhub/LTR_Finder | | [13] |
| Maker2 (includes SNAP) | 2.31.10 | https://www.yandell-lab.org/software/maker_install.html | | [14] |
| MITOS | | http://mitos2.bioinf.uni-leipzig.de/index.py | | [15] |
| MUMmer | | https://github.com/mummer4/mummer | | [16] |
| Pilon | 1.22 | https://github.com/broadinstitute/pilon/wiki | -Xmx160G -- --threads 16 --verbose --changes --fix bases --chunksize 8000000 --diploid | [17] |
| QTLseqr | v0.7.5.2 | https://github.com/bmansfeld/QTLseqr | | [18] |
| RepeatMasker | 4.1.0 | https://github.com/rmhubley/RepeatMasker | -cutoff 300<br>-xsmall<br>-xm<br>-lcambig | [19] |
| RepeatModeler (includes RepeatClassifier) | 2.0.1 | https://github.com/Dfam-consortium/RepeatModeler | For RepeatClassifier<br>-engine ncbi | [20] |
| Rqtl | 1.46.2 | https://github.com/kbroman/qtl | | [21] |
| Samtools | 0.1.19-96b5f2294a | http://samtools.sourceforge.net/ | | [22] |

| | | | | |
|---|---|---|---|---|
| SeqKit | 0.16.1 | https://bioinf.shenwei.me/seqkit/ | | [23] |
| Skewer | v0.2.2 | https://github.com/relipmoc/skewer | -n -Q 20 -l 51 -t 32 -m pe | [24] |
| SnpEff | 5.0.1 | https://pcingola.github.io/SnpEff/download/ | | [25] |
| SnpSift | 4.3.1 | https://pcingola.github.io/SnpEff/download/ | | [26] |
| TransposonPSI | 08222010 | http://transposonpsi.sourceforge.net/ | | [27] |
| Trinity | 2.9.1 | https://github.com/trinityrnaseq/trinityrnaseq | | [28] |
| USEARCH | 11.0.667 | https://www.drive5.com/usearch/download.html | -id 0.9<br>-sort length | [29] |
| VCFtools | 0.1.16 | https://github.com/vcftools/vcftools | | [2] |


1. Tang H, Zhang X, Miao C, Zhang J, Ming R, Schnable JC, Schnable PS, Lyons E, Lu J: **ALLMAPS: robust scaffold ordering based on multiple maps.** *Genome Biol* 2015, **16:**3.
2. Danecek P, Auton A, Abecasis G, Albers CA, Banks E, DePristo MA, Handsaker RE, Lunter G, Marth GT, Sherry ST, et al: **The variant call format and VCFtools.** *Bioinformatics* 2011, **27:**2156-2158.
3. Camacho C, Coulouris G, Avagyan V, Ma N, Papadopoulos J, Bealer K, Madden TL: **BLAST+: architecture and applications.** *BMC Bioinformatics* 2009, **10:**421.
4. Simão FA, Waterhouse RM, Ioannidis P, Kriventseva EV, Zdobnov EM: **BUSCO: assessing genome assembly and annotation completeness with single-copy orthologs.** *Bioinformatics* 2015, **31:**3210-3212.
5. Li H, Durbin R: **Fast and accurate short read alignment with Burrows-Wheeler transform.** *Bioinformatics* 2009, **25:**1754-1760.
6. Koren S, Walenz BP, Berlin K, Miller JR, Bergman NH, Phillippy AM: **Canu: scalable and accurate long-read assembly via adaptive k-mer weighting and repeat separation.** *Genome Res* 2017, **27:**722-736.
7. Kurtz S, Phillippy A, Delcher AL, Smoot M, Shumway M, Antonescu C, Salzberg SL: **Versatile and open software for comparing large genomes.** *Genome Biol* 2004, **5:**R12.
8. Patel RK, Jain M: **NGS QC Toolkit: a toolkit for quality control of next generation sequencing data.** *PLoS One* 2012, **7:**e30619.

9. Pracana R, Priyam A, Levantis I, Nichols RA, Wurm Y: **The fire ant social chromosome supergene variant Sb shows low diversity but high divergence from SB.** *Mol Ecol* 2017, **26:**2864-2879.
10. Van der Auwera GA, Carneiro MO, Hartl C, Poplin R, Del Angel G, Levy-Moonshine A, Jordan T, Shakir K, Roazen D, Thibault J, et al: **From FastQ data to high confidence variant calls: the Genome Analysis Toolkit best practices pipeline.** *Curr Protoc Bioinformatics* 2013, **43:**11 10 11-11 10 33.
11. Lomsadze A, Ter-Hovhannisyan V, Chernoff YO, Borodovsky M: **Gene identification in novel eukaryotic genomes by self-training algorithm.** *Nucleic Acids Res* 2005, **33:**6494-6506.
12. Gremme G, Steinbiss S, Kurtz S: **GenomeTools: a comprehensive software library for efficient processing of structured genome annotations.** *IEEE/ACM Trans Comput Biol Bioinform* 2013, **10:**645-656.
13. Xu Z, Wang H: **LTR_FINDER: an efficient tool for the prediction of full-length LTR retrotransposons.** *Nucleic Acids Res* 2007, **35:**W265-268.
14. Holt C, Yandell M: **MAKER2: an annotation pipeline and genome-database management tool for second-generation genome projects.** *BMC Bioinformatics* 2011, **12:**491.
15. Donath A, Juhling F, Al-Arab M, Bernhart SH, Reinhardt F, Stadler PF, Middendorf M, Bernt M: **Improved annotation of protein-coding genes boundaries in metazoan mitochondrial genomes.** *Nucleic Acids Res* 2019, **47:**10543-10552.
16. Krzywinski M, Schein J, Birol I, Connors J, Gascoyne R, Horsman D, Jones SJ, Marra MA: **Circos: an information aesthetic for comparative genomics.** *Genome Res* 2009, **19:**1639-1645.
17. Walker BJ, Abeel T, Shea T, Priest M, Abouelliel A, Sakthikumar S, Cuomo CA, Zeng Q, Wortman J, Young SK, Earl AM: **Pilon: an integrated tool for comprehensive microbial variant detection and genome assembly improvement.** *PLoS One* 2014, **9:**e112963.
18. Mansfeld BN, Grumet R: **QTLseqr: An R Package for Bulk Segregant Analysis with Next-Generation Sequencing.** *Plant Genome* 2018, **11**.
19. **RepeatMasker-4.0** [www.repeatmasker.org]
20. **RepeatModeler Open-1.0** [http://www.repeatmasker.org]
21. Broman KW, Wu H, Sen S, Churchill GA: **R/qtl: QTL mapping in experimental crosses.** *Bioinformatics* 2003, **19:**889-890.
22. Li H, Handsaker B, Wysoker A, Fennell T, Ruan J, Homer N, Marth G, Abecasis G, Durbin R, Genome Project Data Processing S: **The Sequence Alignment/Map format and SAMtools.** *Bioinformatics* 2009, **25:**2078-2079.
23. Shen W, Le S, Li Y, Hu F: **SeqKit: A Cross-Platform and Ultrafast Toolkit for FASTA/Q File Manipulation.** *PLoS One* 2016, **11:**e0163962.
24. Jiang H, Lei R, Ding SW, Zhu S: **Skewer: a fast and accurate adapter trimmer for next-generation sequencing paired-end reads.** *BMC Bioinformatics* 2014, **15:**182.
25. Cingolani P, Platts A, Wang le L, Coon M, Nguyen T, Wang L, Land SJ, Lu X, Ruden DM: **A program for annotating and predicting the effects of single nucleotide polymorphisms, SnpEff: SNPs in the genome of Drosophila melanogaster strain w1118; iso-2; iso-3.** *Fly (Austin)* 2012, **6:**80-92.
26. Cingolani P, Patel VM, Coon M, Nguyen T, Land SJ, Ruden DM, Lu X: **Using Drosophila melanogaster as a Model for Genotoxic Chemical Mutational Studies with a New Program, SnpSift.** *Front Genet* 2012, **3:**35.
27. **TransposonPSI: An Application of PSI-Blast to Mine (Retro-)Transposon ORF Homologies** [https://transposonpsi.sourceforge.net/]
28. Grabherr MG, Haas BJ, Yassour M, Levin JZ, Thompson DA, Amit I, Adiconis X, Fan L, Raychowdhury R, Zeng Q, et al: **Full-length transcriptome assembly from RNA-Seq data without a reference genome.** *Nat Biotechnol* 2011, **29:**644-652.
29. Edgar RC: **Search and clustering orders of magnitude faster than BLAST.** *Bioinformatics* 2010, **26:**2460-2461.
