## Supplementary material for "The contribution of an X chromosome QTL to non-Mendelian inheritance and unequal chromosomal segregation in *A. freiburgense*": Peer review report

### Peer Reviewer Report ©

Peer Premier Inc
(all rights reserved)

**Manuscript Number** PPP-2023-6-2

Digitally signed by Peer Premier
Date: 2023.08.24 14:21:44 -04'00'

**Title:**

The contribution of an X chromosome QTL to non-Mendelian inheritance and unequal chromosomal segregation in A. freiburgense

Network Analysis (peer reviewer subject areas; created by vosviewer.com):

Sex determination, QTL, NOT Fish

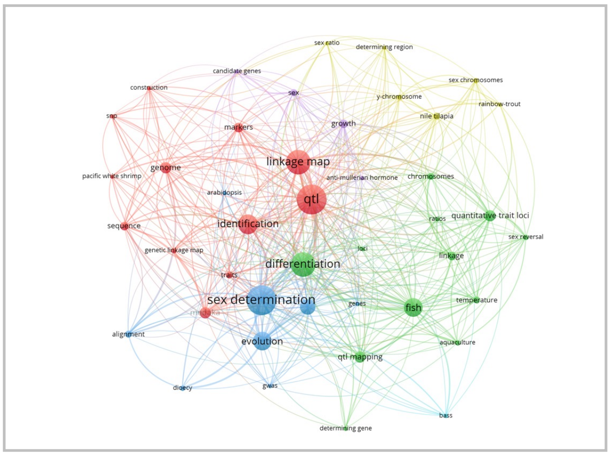


315 Authors (Anonymized)

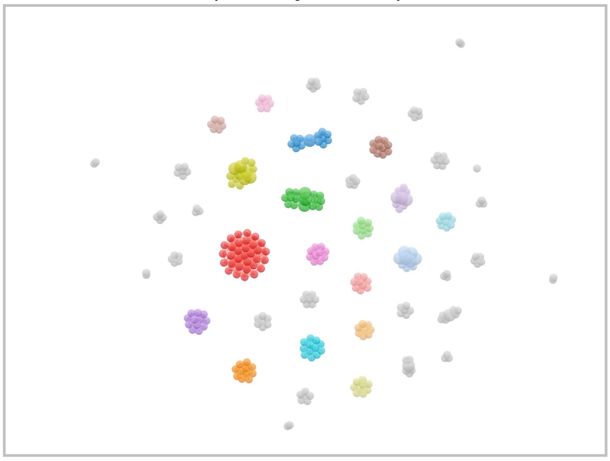

Level/rank of peer reviewers: Senior Research Fellow; Professor; Senior Lecturer

| | Reviewer Average | Coefficient of Variation (%) |
|---|---|---|
| Quality: | 8.6 | 9 |
| Novelty: | 8.9 | 9 |
| Importance/Relevance: | 8.5 | 12 |
| Presentation/Clarity: | 8.7 | 5 |

**Strong Points:** substantial effort and interesting biology described this manuscript makes a significant contribution to the field.: a beautiful illustration of how modern genomics can be applied to a non-model organism; a substantial piece of work that bring together multiple genetic techniques to explore a particular pattern of segregation and organelle partitioning within a species of nematodes.

**Weak Points:** While the authors are able to narrow down a list of candidates they do not discuss or list the annotated functions of these genes; I feel that many of the discussion points are out of sync with the results presented; Expansion of the discussion is needed to place these results within the broader literature and future applications.

The coefficient of variation is a reflection of reviewer cohesiveness. The closer the value to zero the more the agreement amongst reviewers.

Disclaimer: This report is the assessment by three independent reviewers using a double-blind review process. The comments and scores have not been altered in any way by Peer Premier. Quantitative and qualitative information provided in this report reflects the sole opinion of these reviewers based on their professional capacity at the time of review and for the specific version of the manuscript that was submitted to Peer Premier. Peer Premier is not responsible for these assessments. The scores and comments are intended to provide authors with high-quality feedback on their manuscript. Consequently, they should not be used to rank or compare reports that have been generated by different peer reviewers. It is Peer Premiers belief that the ultimate impact of the paper will be determined by the research community.

#### Title

(1=Disagree - 10=Agree)

**Accurately reflects the conclusions of the manuscript.**

| | | | |
|---|---|---|---|
| 10.0 | 8.0 | 9.0 | **9.0** |

**Is sufficiently detailed enough to get the point across.**

| | | | |
|---|---|---|---|
| 9.0 | 7.0 | 9.0 | **8.3** |

**Does not overstate the conclusions of the paper.**

| | | | |
|---|---|---|---|
| 9.0 | 9.0 | 9.0 | **9.0** |

##### Comments on the title.

Reviewer #1

appropriate
Author Response:
NA

Reviewer #2

I would focus it on the non-Mendelian inheritance of the X chromosome rather than sex ratio
Author Response:
We followed this recommendation

Reviewer #3

Concise and accurate title.
Author Response:
Another reviewer asked us to change the title to: The contribution of an X chromosome QTL to

#### Abstract

(1=Disagree - 10=Agree)

**The abstract is clear and succinct.**

| | | | |
|---|---|---|---|
| 9.0 | 8.0 | 9.0 | **8.7** |

**The abstract draws accurate and meaningful conclusions.**

| | | | |
|---|---|---|---|
| 9.0 | 8.0 | 8.0 | **8.3** |

**The abstract accurately reflects the main points of the paper.**

| | | | |
|---|---|---|---|
| 9.0 | 8.0 | 8.0 | **8.3** |

##### Comments on the Abstract.

Reviewer #1

Overall the abstract is clear and well-written although the last sentence "We found a QTL on the X chromosome associated with the unusual spermatogenesis in A. freiburgense, providing a mechanistic basis for asymmetric partitioning of organelles during the male meiotic division."

Reviewer #2

Same comment as on the title, put more focus on the non-Mendelian inheritance of the X chromosome
Author Response:

Reviewer #3

The abstract is quite brief and does not really cover the full scope of the manuscript. Adding a few more sentences to expand on the manuscripts context and results would add value here
Author Response:

#### Material and Methods

(1=Mostly Incomplete - 10=Mostly Complete)

**Description of the material and methods.**

| | | | |
|---|---|---|---|
| 9.0 | 10.0 | 9.0 | **9.3** |

**Sources of materials (references, manufacturer, etc).**

| | | | |
|---|---|---|---|
| 9.0 | 9.0 | 9.0 | **9.0** |

**Details that would allow replication of this study .**

| | | | |
|---|---|---|---|
| 9.0 | 8.0 | 9.0 | **8.7** |

**Ethics/animal approval with appropriate protocol identifiers have been included?**

| Reviewer #1 | Reviewer #2 | Reviewer #3 |
|---|---|---|
| Not Required | Not Required | Not Required |

##### Comments on the Material and Methods.

Reviewer #1

Very detailed, clear and well-structured
Author Response:
NA

Reviewer #2

clear
Author Response:
NA

Reviewer #3

Methods were comprehensive and would allow the replication of the study.
Author Response:
NA

#### Results

##### Model

| | | Neccessary? | How well described? |
|---|---|---|---|
| **Please indicate if the source of the model is necessary for proper evaluation of the results or to reproduce the experiment.** | R1 | Yes | 9.0 |
| | R2 | Yes | 10.0 |
| | R3 | Yes | 9.0 |
| **Please indicate if the sex and age of the model is necessary for proper evaluation of the results or to reproduce the experiment.** | R1 | Yes | 8.0 |
| | R2 | Yes | 9.0 |
| | R3 | Yes | 9.0 |
| **Please indicate if the number of replicates per experiment is necessary for proper evaluation of the results or to reproduce the experiment.** | R1 | Yes | 9.0 |
| | R2 | No | |
| | R3 | Yes | 9.0 |

Peer Premier Certified

#### Model (cont'd)

| | | Neccessary? | How well described? |
|---|---|---|---|
| Please indicate if the strain or line description (eg. transgenic, mutant, reporter, etc) is necessary for proper evaluation of the results or to reproduce the experiment. | R1 | Yes | 9.0 |
| | R2 | Yes | 9.0 |
| | R3 | Yes | 9.0 |
| Please indicate if controls are necessary for proper evaluation of the results or to reproduce the experiment. | R1 | No | |
| | R2 | No | |
| | R3 | Yes | 8.0 |

**Based on the information above, rate your overall satisfaction with the description of the model used for this particular point,**

9.0 | 9.0 | 9.0 | **9.0**

**Please provide comments on the model(s) used for this study (suitability, alternatives, etc).**

Reviewer #1

unique biology, no alternative
Author Response:
NA

Reviewer #2

add a bit information on the role of organelles in spermatogenesis
Author Response:
More information was added to the Introduction, lines 95-102

Reviewer #3

Nematodes are an ideal model organisim to undertake such studies due to their short generation intevals, relatively easy rearing within lab conditions and ease of pairing individuals to breed.
Mulitple lines were utilised in this experiment that produced many crosses to be investigated

#### Techniques

**Using the slider below, are the techniques (eg., Western blot, xenograph, luciferase reporter, X-ray, etc) appropriate to address the point in question? (1=Not appropriate - 10=very appropriate)**

9.0 | 10.0 | 9.0 | **9.3**

**Please provide comments on the techniques and identify limitations and improvements.**

Reviewer #1

QTL methods used widely established and appropriate
Author Response:
NA

Reviewer #2

no comments
Author Response:
NA

Reviewer #3

A large array of techniques was used within this manuscript. Each were appropriate in their own right. The authors also produced a genome, which then formed a common anchor point between QTL mapping studies and the other methods utilised in the

#### Experimental Design (1=Disagree - 10=Agree)

**These are unique experiments, not duplicates of previous published experiments.**

9.0 | 10.0 | 9.0 | **9.3**

**The results are new. That is they are not simply confirming known results.**

9.0 | 10.0 | 9.0 | **9.3**

**The experimental design is adequate for the point the authors are trying to make.**

9.0 | 10.0 | 9.0 | **9.3**

**Comments on the Experimental Design.**

Reviewer #1

good overall
Author Response:
NA

Reviewer #2

beautiful experimental design
Author Response:
NA

Reviewer #3

This study is unique in the way that it employs QTL mapping and other techniques to explore not only the sex chromosomes, but also the partitioning of cellular organelles within germ cell production.

#### Novelty

**How novel are the models, techniques, technologies and experiments that the authors use to use address this point? (1=Existing Technology - 10=New Innovation)**

8.0 | 10.0 | 8.0 | **8.7**

**How does this research build on existing research? (1=Does not advance - 10=Significantly advances)**

8.0 | 9.0 | 9.0 | **8.7**

**Assess the overall novelty of the manuscript based on the methods and the results obtained. (1=Lacks novelty - 10=Very novel)**

7.0 | 10.0 | 9.0 | **8.7**

**Comments on the novelty and potential innovation.**

Reviewer #1

described a new not previously observed phenotype, generated a previously unavailable reference genome and where able to map the phenotype to a X-linked region
Author Response:

Reviewer #2

opens up this new model system for investigating unequal chromosome segregation
Author Response:
NA

Reviewer #3

The techniques themselves are commonly used to address a range of scientific questions, however, the implementation of this set of techniques to the research question here is unique.

Peer Premier Certified

#### Statistics

Rate your overall satisfaction with the statistics for this manuscript. (1=Dissatisfied - 10=Satisfied)

9.0 | 9.0 | 9.0 | **9.0**

Rate your overall satisfaction with the number of biological replicates. (1=Dissatisfied - 10=Satisfied)

9.0 | 8.0 | 9.0 | **8.7**

Is the data supported by the appropriate statistics?

| | R1 | R2 | R3 |
|---|---|---|---|
| Statistics are not required. | ☐ | ☐ | ☐ |
| I am unable to accurately evaluate the statistics used. | ☐ | ☐ | ☐ |
| Replicates not provided when they should be. | ☐ | ☐ | ☐ |
| Multiple reps provided but not statistically evaluated. | ☐ | ☐ | ☐ |
| No statistics were included when they should have been. | ☐ | ☐ | ☐ |
| Some statistics included or stats included but of the wrong type or stats included but indication of the statistic that were used or the parameters of the stats not included. | ☐ | ☐ | ☐ |
| Statistics are included and may be of the right type but information is missing. | ☐ | ☐ | ☐ |
| Statistics are included, of the right type with appropriate information included. | ✔ | ✔ | ✔ |

Other

Comment on the statistics.

Reviewer #1

statical methods appropriate and clearly explained
Author Response:
NA

Reviewer #2

appropriate QTL analysis
Author Response:
NA

Reviewer #3

The statistics employed throughout the manuscript are specific to each analysis and were appropriately documented, presented and interpreted.
Author Response:

#### Controls (1=None of the time - 10=All of the time)

Are negative controls provided?

5.0 | 5.0 | 5.0 | **5.0**

Are positive controls provided?

5.0 | 5.0 | 5.0 | **5.0**

Are the controls used appropriate?

10.0 | 9.0 | 9.0 | **9.0**

Do the authors justify their use of controls?

10.0 | 9.0 | 9.0 | **9.3**

Comment on the controls.

Reviewer #1

No need for controls in the experimental design used
Author Response:
NA

Reviewer #2

there are no real negative and positive controls to this study
Author Response:
NA

Reviewer #3

Within this study, the authors produced 100 lines across many different crosses to explore patterns of inheritance and expression between the major sexual morphs of nematodes. This enabled them to set various scenarios to investigate the percentage

#### Evidence (1=Poor - 10=Excellent)

What is your overall evaluation of the readability of the data presented? (clear labels, units, , font size, etc)

9.0 | 8.0 | 9.0 | **8.7**

What is your overall evaluation of the organization of the data?

9.0 | 9.0 | 9.0 | **9.0**

What is your overall evaluation of the quality of the data presented?

9.0 | 9.0 | 9.0 | **9.0**

What is your overall evaluation of the presentation of the data?

9.0 | 8.0 | 9.0 | **8.7**

Evaluation is...? (1=missing - 10=complete)

9.0 | 9.0 | 9.0 | **9.0**

Peer Premier Certified

#### Evidence Cont'd

**Please provide specific comments about each of the points that the authors are trying to make. For example you may agree with their conclusions about some experiments, but not others, so please elaborate.**

Reviewer #1

The power of the QTL approach to fine-map the trait are limited but the authors acknowledge this and other mapping approaches are not feasible
Author Response:

Reviewer #2

results are appropriately interpreted
Author Response:
NA

Reviewer #3

The authors have done a great job of being thorough with this manuscript, and not overstepping conclusions. They point out where their power was insufficient rather than speculating conclusions and outline future methodology that may help address remaining

#### Depth

**Your overall satisfaction with the number of experiments or approaches used to address their topic (1=Very Dissatisfied - 10=Very Satisfied)**

9.0 | 8.0 | 9.0 | **8.7**

**Please provide specific comments about each of the points that the authors are trying to make regarding the depth of the research. For example, some questions may be fully addressed, but other questions may only have preliminary data, so please elaborate.**

Reviewer #1

Overall, the manuscript describes an interesting polymorphic sex determining phenotype, generates a reference genome that will substantially aid further work on this interesting nematode model, show that an X-linked factor is probably involved in regulating the

Reviewer #2

no comments
Author Response:
NA

Reviewer #3

This manuscript covers a wide range of experiments and a substantial amount of analysis which explores their questions in-depth. Often studies like this explore only on or the other between transcriptome analysis and QTL, however this manuscript reports both.

#### Importance and Relevance

**Rate the overall importance of the research to the main objective of the paper.(1=Not Important - 10=Very Important)**

7.0 | 8.0 | 9.0 | **8.0**

#### Importance and Relevance Cont'd

**Rate the overall importance of the research to the specific field of study. (1=not important - 10=very important)**

8.0 | 10.0 | 9.0 | **9.0**

**Rate the overall importance of the research to the scientific field in general. (1=not important - 10=very important)**

7.0 | 9.0 | 9.0 | **8.3**

**Please provide specific comments regarding importance and relevance. For example, some experiments may be more important than others, so please elaborate.**

Reviewer #1

Sex determining mechanisms are incredibly diverse, and in many cases the molecular mechanisms remain poorly understood, with the exception of just a handful of model systems. Even less is know about more unusual sex determining systems that differ from

Reviewer #2

see my concerns below and my report with suggestions to frame the study more appropriately
Author Response:

Reviewer #3

Chromosome evolution, especially of sex chromosomes and sex determining systems is a fundamental part of biology. This study reports a detailed examination of spermatogenesis and the segregation of males and females within many lines of

#### Concerns

**Do you have any concerns about this manuscript that have not been addressed with the previous questions? (1=Major Concerns - 10=No Concerns)**

8.0 | 9.0 | 9.0 | **8.7**

**Please provide specific comments regarding any concerns you may have.**

Reviewer #1

The introduction of the manuscript, while giving a nice introduction into the system is quite concise to a point that the last paragraph where the author summarize the approach and results of the study is hard to follow for someone not intimately familiar with the

Reviewer #2

This is a wonderful study into the genetics of an extraordinary reproductive system. We need many more of such studies to better understand the evolution of unusual reproductive modes and non-Mendelian systems of sex determination. It is also a

Reviewer #3

No real concerns on this manuscript or work.
Author Response:
Thank you very much for your comments.

Peer Premier Certified

#### Discussion (1=Disagree - 10=Agree)

**The discussion avoids unnecessarily repeating the results.**

| | | | |
|---|---|---|---|
| 9.0 | 9.0 | 9.0 | **9.0** |

**The main conclusions are justified by the results presented.**

| | | | |
|---|---|---|---|
| 9.0 | 5.0 | 9.0 | **7.7** |

**The discussion puts the results into context with the current understanding in the field.**

| | | | |
|---|---|---|---|
| 8.0 | 4.0 | 8.0 | **6.7** |

**Limitations have been discussed.**

| | | | |
|---|---|---|---|
| 8.0 | 8.0 | 8.0 | **8.0** |

**Authors acknowledge obvious unanswered questions.**

| | | | |
|---|---|---|---|
| 8.0 | 7.0 | 8.0 | **7.7** |

**Please provide comments or concerns you may have regarding the discussion.**

Reviewer #1

One thing that is missing for me, especially in the discussion, but also in the discussion is for the authors to place the system and their findings into the wider context of unusual sex determination systems that involve DNA/sperm elimination and of sex determination in

Reviewer #2

See my concerns Discussion, page 23, 1st par.: this paragraph is a bit out of sink with the subtitle [see also conceptual comments above] Page 24: Transgressive phenotypes: this is also deferring from the main findings. The interesting discussion point

Reviewer #3

The discussion is quite concise and could benefit from some elaborations in some cases. Most of this is expanding out their findings within the literature and drawing comparisons between similar studies either in nematodes or a wider array of species. Explicitly stating

#### References (1=Disagree - 10=Agree)

**Authors include the most up to date and relevant (primary research) references.**

| | | | |
|---|---|---|---|
| 7.0 | 8.0 | 9.0 | **8.0** |

**The references are sufficiently extensive.**

| | | | |
|---|---|---|---|
| 7.0 | 8.0 | 9.0 | **8.0** |

**Please provide comments or concerns you may have regarding the references.**

Reviewer #1

Very limited number of reference but sufficient
Author Response:
NA

Reviewer #2

See my concerns Include more refences to extraordinary chromosomal inheritance systems and studies
Author Response:

Reviewer #3

The authors present an expansive list of relevant references.
Author Response:
NA

Peer Premier Certified

### Major and Minor Points

#### Strong Points of the paper.

Reviewer #1

Sex determining mechanisms are incredibly diverse, and in many cases the molecular mechanisms remain poorly understood, with the exception of just a handful of model systems. Even less is know about more unusual sex determining systems that differ from the standard sex chromosome systems most of us are familiar with. One type of unusual sex determination which has repeatedly evolved across diverse organisms is sex determination through DNA elimination. Such systems have evolved in rodents, several fly species as well as nematodes but remain poorly understood especially at the molecular level. One challenge in identifying the molecular underpinnings of DNA elimination is that the phenomena is often an integral part of the life cycle and therefore within-population polymorphisms are rare. In this manuscript the authors report such polymorphism in a species of nematode where males are XO but produce an excess of X-bearing sperm, and therefore an excess of daughters, as result of the elimination of nullo-X sperm. They found that the F1 offspring from a cross between two inbred lines produced significantly more male offspring and are able to attribute

Reviewer #2

This is a wonderful study into the genetics of an extraordinary reproductive system. We need many more of such studies to better understand the evolution of unusual reproductive modes and non-Mendelian systems of sex determination. It is also a beautiful illustration of how modern genomics (e.g. chromosome-level assembly) can be applied to a non-model organism. The study is well-designed, the methods clearly described and the results well presented.

Reviewer #3

The manuscript is a substantial piece of work that bring together multiple genetic techniques to explore a particular pattern of segregation and organelle partitioning within a species of nematodes.

#### Weak Points of the paper.

Reviewer #1

The introduction of the manuscript, while giving a nice introduction into the system is quite concise to a point that the last paragraph where the author summarize the approach and results of the study is hard to follow for someone not intimately familiar with the methods and study system. For example, it was unclear to me what the authors meant by "transgressive phenotype" until much further into the manuscript. I would suggest maybe adding one extra paragraph of introduction explaining this phenotype and how it was derived. While the authors are able to narrow down a list of candidates they do not discuss or list the annotated functions of these genes. Are there any genes with functions that could indicate that they might be promising candidates, such as genes involved in meiosis or spindle behaviour? One thing that is missing for me, especially in the discussion, but also in the discussion is for the authors to place the system and their findings into the wider context of unusual sex determination systems that involve DNA/sperm elimination and of sex determination in nematodes. Doing so would likely make the manuscript more appealing to a wider audience. Maybe also

Reviewer #2

I feel that many of the discussion points are out of sink with the results presented. This already starts with the first paragraph of the Introduction that focuses on sex ratio evolution, but this system is not very appropriate to test sex allocation theory. Moreover, the authors do not come back to sex ratio evolution in the Discussion. The same holds for transgressive phenotypes, which makes up a major part of the Discussion. Transgressive phenotypes is not the most important finding of this study, and there is no evidence for hybrid formation.

Reviewer #3

Expansion of the abstract is needed for more context. Expansion of the discussion is needed to place these results within the broader literature and future applications.

#### Major Points of concern not addressed above.

Reviewer #1

Reviewer #2

See concerns above I copy here my reviewer report that contains most of the points raise above, but also some additional ones. The authors present a study into the genetic basis of sex determination in the nematode Auanema freiburgense. This nematode has an extraordinary reproductive mode, consisting of hermaphroditic females, XX females and XO males. Males produce sperm with and without an X chromosome, but the nullo-X sperm has reduced viability, causing non-Mendelian inheritance of the X through males resulting in biased sex ratios. Viability of the nullo-X sperm depends on the asymmetry of the spermatogonial spindle and the associated segregation of organelles. By making crosses between lines that differ in the degree of nullo-X sperm viability, the authors find a QTL underlying the polarized segregation of the X chromosome during spermatogenesis. This is a wonderful study into the genetics of an extraordinary reproductive system. We need many more of such studies to better understand the evolution of unusual reproductive modes and non-Mendelian systems of sex determination. It is also a beautiful illustration of how modern

Reviewer #3

NA

#### Minor Points of concern not addressed above.

Reviewer #1

"In the ~800 Mb mapped QTL region there are 123 predicted protein-coding genes, and 75 of the candidate protein-coding genes have at least one missense mutation." Is this a typo? The QTL is presumably 0.8 Mb given the whole genome is significantly less than 800Mb

Reviewer #2

a few textual comments

Reviewer #3

Na

Peer Premier Certified
